# Dual inhibition of coronavirus M^pro^ and PL^pro^ enzymes by phenothiazines and their antiviral activity

**DOI:** 10.1101/2023.09.11.557219

**Authors:** Katrina Forrestall, Eric S. Pringle, Dane Sands, Brett A. Duguay, Brett Farewell, Tekeleselassie Woldemariam, Darryl Falzarano, Ian Pottie, Craig McCormick, Sultan Darvesh

## Abstract

Coronavirus (CoV) replication requires efficient cleavage of viral polyproteins into an array of non-structural proteins involved in viral replication, organelle formation, viral RNA synthesis, and host shutoff. Human CoVs (HCoVs) encode two viral cysteine proteases, main protease (M^pro^) and papain-like protease (PL^pro^), that mediate polyprotein cleavage. Using a structure-guided approach, a phenothiazine urea derivative that inhibits both SARS-CoV-2 M^pro^ and PL^pro^ protease activity *in vitro* was identified. *In silico* docking studies also predicted binding of the phenothiazine to the active sites of M^pro^ and PL^pro^ from distantly related alphacoronavirus, HCoV-229E (229E) and the betacoronavirus, HCoV-OC43 (OC43). The lead phenothiazine urea derivative displayed broad antiviral activity against all three HCoVs tested in cell culture infection models. It was further demonstrated that the compound inhibited 229E and OC43 at an early stage of viral replication, with diminished formation of viral replication organelles and the RNAs that are made within them, as expected following viral protease inhibition. These observations suggest that the phenothiazine urea derivative inhibits viral replication and may broadly inhibit proteases of diverse coronaviruses.

**Graphical Abstract:** 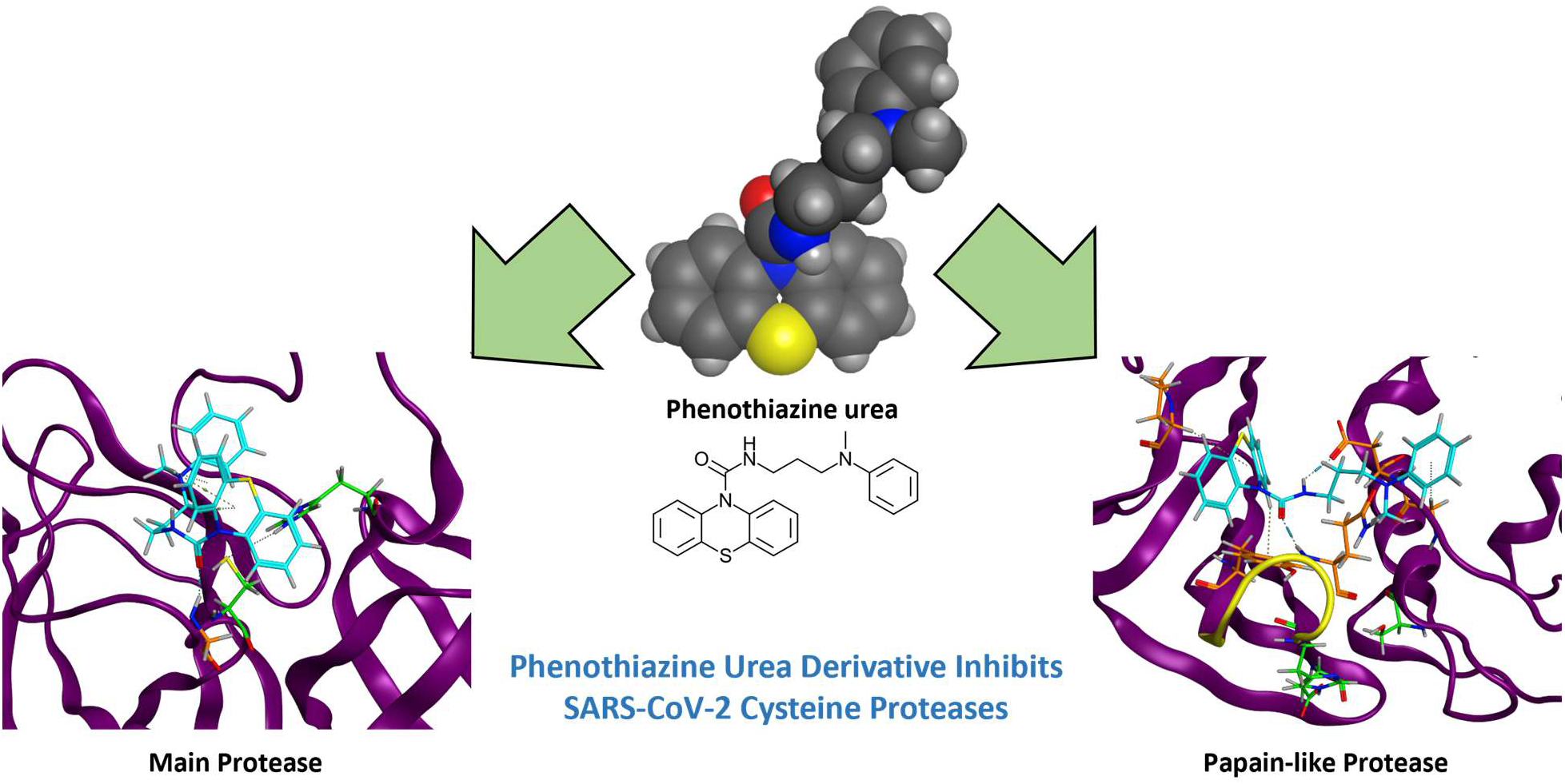

**Highlights:**

- Coronavirus cysteine proteases M^pro^ and PL^pro^ are targets for novel antiviral agents
- Phenothiazine ureas inhibit SARS-CoV-2 M^pro^ and PL^pro^ protease activity
- Some phenothiazine ureas inhibit replication of diverse coronaviruses with minimal cytotoxicity
- Phenothiazine ureas inhibit early stages of coronavirus replication consistent with failure of viral polyprotein cleavage

## 1.0 INTRODUCTION

Coronaviruses (CoVs) are enveloped viruses with ∼30 kilobase positive-sense, single-stranded RNA ((+)ssRNA) genomes [Zhou et al., 2021]. Seven human infectious CoVs (HCoVs) are divided into two genera, the alphacoronaviruses (HCoV-NL63 and HCoV-229E) and the betacoronaviruses (HCoV-OC43, HCoV-HKU1, MERS-CoV, SARS-CoV and SARS-CoV-2). Upon entry into host cells, the (+)ssRNA CoV genome is translated into polyproteins that are cleaved by the viral cysteine proteases, main protease (M^pro^ or 3CL^pro^) and papain-like protease (PL^pro^), into 16 non-structural proteins that play critical roles in viral replication [Ullrich and Nitsche 2022; Tan et al., 2022]. M^pro^ and PL^pro^ enzymes are also responsible for self-cleavage into their respective NSPs, NSP5 and NSP3. Beyond the processing of viral polyproteins, PL^pro^ has additional roles in suppressing host immune defences by acting as a ubiquitin and ubiquitin-like interferon-stimulated gene 15 (ISG15) deconjugating enzyme [Shin et al., 2021; Ullrich and Nitsche, 2022; Tan et al., 2022]. Other studies have shown M^pro^ activity against host proteins involved in immune response, such as galectin-8 [Pablos et al., 2021; Chen et al., 2023]. The proteolytic and immune modulating functions of these enzymes make M^pro^ and PL^pro^ attractive targets for potential antiviral drug intervention [Owen et al., 2021; Cao et al., 2023; Puhl et al., 2023]

The active site of M^pro^ contains a catalytic dyad (C145 and H41) [Zhang et al., 2020a, b] (Figure 1A, C), while the active site of PL^pro^ contains a catalytic triad (C111, H273, and D287) [Klemm et al., 2020; Osipiuk et al., 2021] (Figure 1B, D). Furthermore, a highly mobile β-loop (BL2 Loop) consisting of amino acids (AAs) 266-272, is located between the active site and S1 binding site of PL^pro^ (Figure 1B, D). The active sites of both proteases have shown high conservation among other alpha- and beta-coronaviruses [Shin et al., 2021; Masood et al., 2020; Ullrich and Nitsche, 2022], presenting an opportunity for the development of broad-spectrum antivirals.

**Figure 1:**
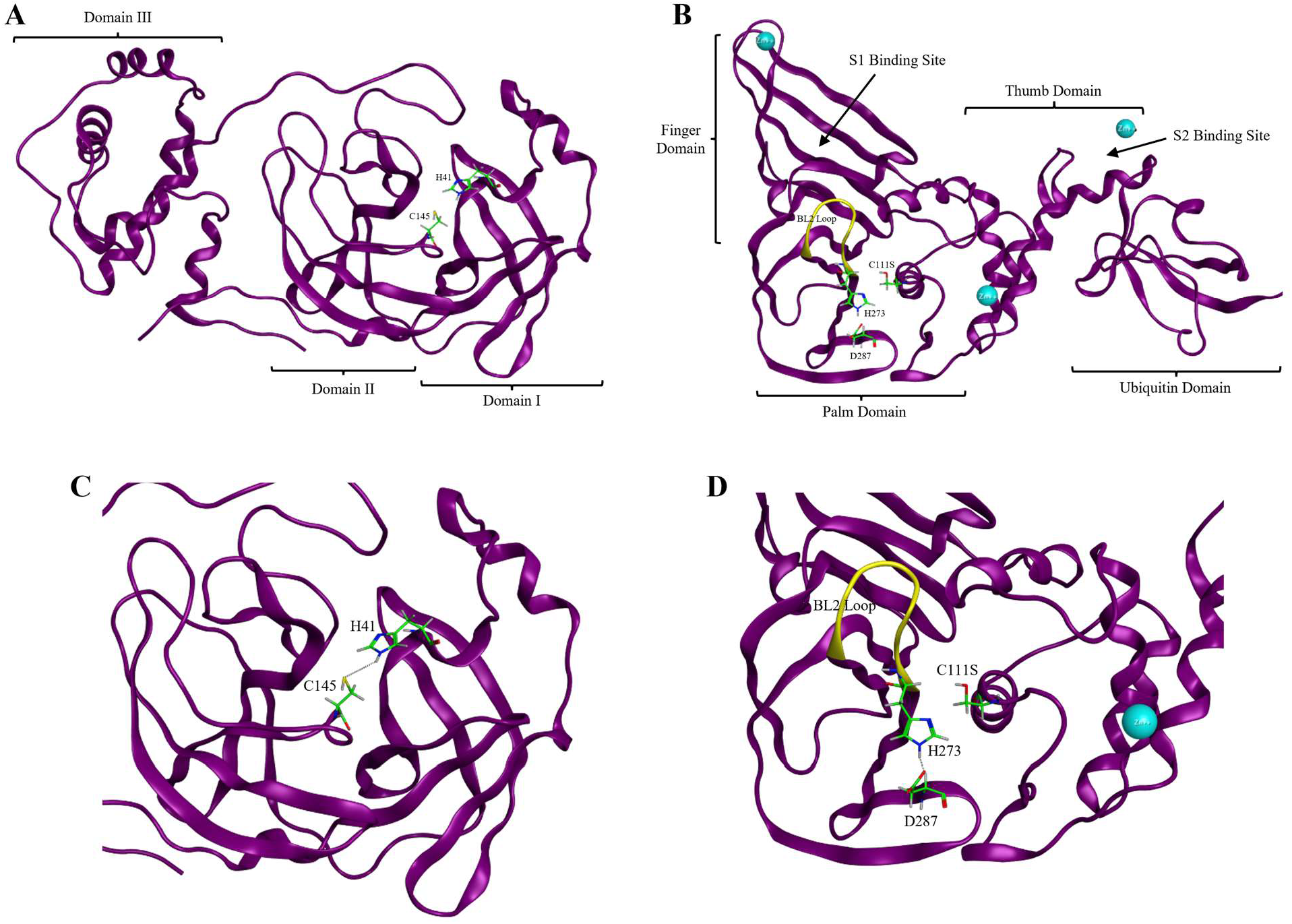
Structural features of SARS-CoV-2 cysteine proteases. Enzyme structure of SARS-CoV-2 main protease (M^pro^; PDB: 7L10) [Zhang et al., 2020a,b] with labelled domains (**A**). Enzyme structure of SARS-CoV-2 papain-like protease (PL^pro^; PDB: 7CJM) [Fu et al., 2021] with labelled domains (**B**). Active site of SARS-CoV-2 M^pro^ with catalytic dyad (H41 and C145) shown as lime green carbon atoms (**C**). Active site of SARS-CoV-2 PL^pro^ with catalytic triad (C111S, H273, and D287) shown as lime green carbon atoms and BL2 Loop (AAs 266-272) shown as yellow ribbon (**D**). Images were generated using Molecular Operating Environment 2022.02 (https://www.chemcomp.com).

Phenothiazines are tricyclic aromatic compounds containing nitrogen and sulfur atom in its center ring. These compounds have been shown to exhibit extraordinary versatility and have great clinical utility for the treatment of a variety of conditions, while also being well tolerated [Baldessarini, 1996; Mitchell, 2006]. Many phenothiazine drugs display reasonable half-lives for daily administration [Baldessarini, 1996; Wishart et al., 2017] without the need for co-formulation with other drugs. In a drug-repurposing study several phenothiazine-containing drugs approved by the FDA for the treatment of other medical conditions have been shown to inhibit spike protein-mediated entry of SARS-CoV-2 into cells [Hashizume et al., 2023]. Previous *in*-*silico* docking studies identified certain phenothiazines with strong predicted binding to the SARS-CoV-2 M^pro^ active site [Forrestall et al., 2021]. Here, we expand on this concept and demonstrate that some novel phenothiazine urea compounds inhibit both M^pro^ and PL^pro^ or SARS-CoV-2 and have broad antiviral action against other coronaviruses.

## 2.0 MATERIALS AND METHODS

### 2.1 Phenothiazine Compounds

Solvents used were purchased from Fisher Scientific (https://www.fishersci.ca), VWR International (https://ca.vwr.com), Sigma-Aldrich (https://www.sigmaaldrich.com), or Oakwood Chemical (https://oakwoodchemical.com). Phenothiazine derivatives **1**-**7** were synthesized and fully characterized as previously described [Darvesh et al, 2010]. Details on the synthetic method and spectroscopic data not previously published can be found in the Supplementary Materials. Synthetic purity was confirmed >99% for all compounds by HPLC.

### 2.2 Enzyme Kinetic Studies

Recombinant M^pro^ (Sigma Aldrich, SAE0172) was resuspended in 10% aqueous glycerol to a concentration of 1 mg/mL. The M^pro^ fluorogenic substrate (Sigma-Aldrich, SAE0180) was solubilized to 1 mg/mL in dimethyl sulfoxide (DMSO). Recombinant PL^pro^ (BPS Biosciences, 100735) was provided in a formulation of 40 mM Tris-HCl buffer (pH 8), 110 mM NaCl, 2.2 mM KCl, 0.04% Tween-20, 3 mM dithiothreitol (DTT), 20% glycerol, and 250 mM imidazole at a concentration of 1.60 mg/mL PL^pro^. This was further diluted to give a stock solution of 1 mg/mL, with 20% glycerol_(aq.)_. The PL^pro^ fluorogenic substrate (BPS Biosciences, 79997) was provided at a concentration of 5 mM (4.2 mg/mL) in DMSO which was further diluted to give a stock solution of 2 mg/mL in DMSO.

To assess enzymatic activity, *N*-2-hydroxyethylpiperazine-*N*-2-ethane sulfonic acid (HEPES) buffer (395.6 µL; 25 mM; pH 7.0) was mixed with 0.4 µL of M^pro^ stock solution or 0.4 µL of PL^pro^ stock solution and 2 µL of their appropriate fluorogenic substrate stock solution in a quartz cuvette of 1 cm path length and measured using a Varian Cary Eclipse Fluorescence spectrophotometer (Varian Inc., Palo Alto CA, USA). Baseline fluorescence was recorded using DMSO in place of fluorogenic substrate. The resulting fluorescence, Relative Fluorescence Units (RFU), was measured until the fluorescence approached an asymptote. Each experiment was repeated in triplicate and values were averaged with standard error. Spectrophotometric settings used for each enzyme were as follows: Excitation at 340 nm (M^pro^) or 330 nm (PL^pro^), emission at 450 nm (M^pro^) or 430 nm (PL^pro^), excitation slit at 2.5 nm, emission slit at 10 nm, emission filter at 360-1100 nm (M^pro^) or *auto* (PL^pro^), excitation filter at 250-395 nm (M^pro^) or *auto* (PL^pro^), with a cycle time of 0.05 min and a total run time of 60 mins.

To evaluate inhibition constants, each phenothiazine compound was dissolved in DMSO at various concentrations (1 µM – 250 µM). Using the same procedure above, each phenothiazine was added to the cuvette prior to the addition of fluorogenic substrate. Rates of hydrolysis were plotted against the concentration of inhibitor and the plot was fit non-linearly to Morrison’s equation as shown in the literature [Hu et al., 2023] using PRISM 9.4.0 (673) (GraphPad Software Inc., San Diego CA, USA). Lineweaver-Burk plots were generated with varying concentrations of compound **5** and a consistent concentration of M^pro^ (30 nM) or PL^pro^ (27 nM) to determine the mode of inhibition. Replotting the calculated slopes from the double reciprocal plots against the solvent concentration gives the inhibitor constant (*K*_i_, M) as the x-intercept.

### 2.3 *In-silico* Docking Studies

To assess the structure activity of phenothiazine derivatives against SARS-CoV-2 M^pro^ and PL^pro^, compounds were docked using a reversible inhibitor docking procedure with Molecular Operating Environment (MOE) 2022.02 (Chemical Computing Group, Montreal, Canada; https://chemcomp.com). The docking procedure consisted of two phases: a blind docking phase in which all ligands were directed to dock anywhere on the enzyme; and a site-directed docking phase in which ligands were directed to dock to the active site of the enzyme. Blind docks were carried out initially to indicate whether a compound is likely to find the active site of the enzyme, followed by site docking to determine possible interactions with enzyme residues and predicted *K*_i_s. Crystal structures of SARS-CoV-2 M^pro^ (Protein Databank code (PDB): 7L10, 1.63 Å) [Zhang et al., 2020a,b] and PL^pro^ (PDB: 7CJM, 3.20 Å) [Fu et al., 2021] were obtained from the Protein Databank (https://rcsb.org) and prepared for molecular docking. Because PL^pro^ could not be crystallized, the catalytic C111 was mutated to S111 which facilitated this process (Figure 1B, D) [Fu et al., 2021]. Though this mutant would be catalytically inactive, it remains useful for screening non-covalent inhibitors *in-silico* [Gao et al., 2021].

The binding energies of the top five returned site-dock poses, key amino acid interactions, and bond lengths were identified and measured within MOE. Docks were completed in triplicate and their binding energies were converted to predicted *K*_i_ values using Gibb’s free energy equation as done previously [Forrestall et al., 2020] and averaged to produce values in Table 1. Further details on the docking procedure can be found in Supplementary Materials.

**Table 1.**
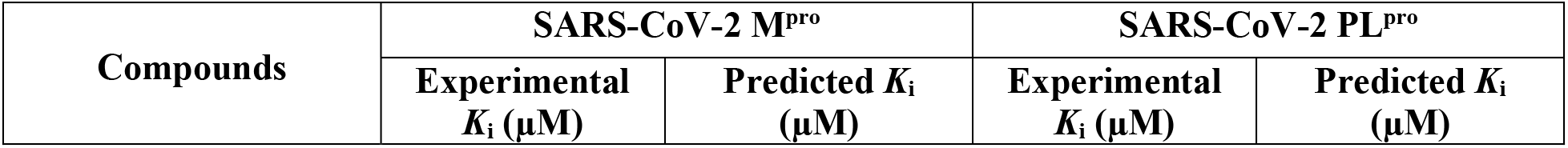
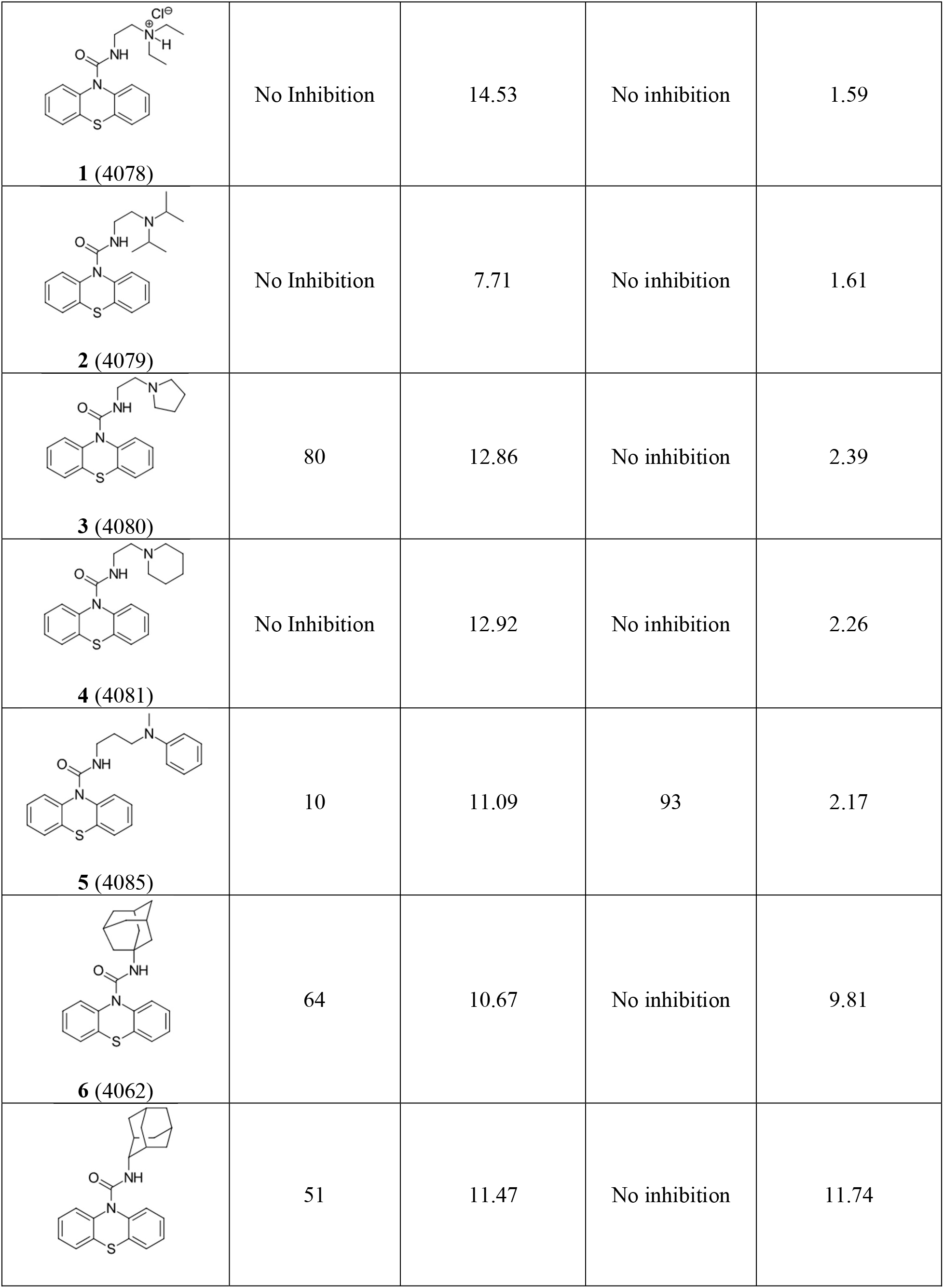
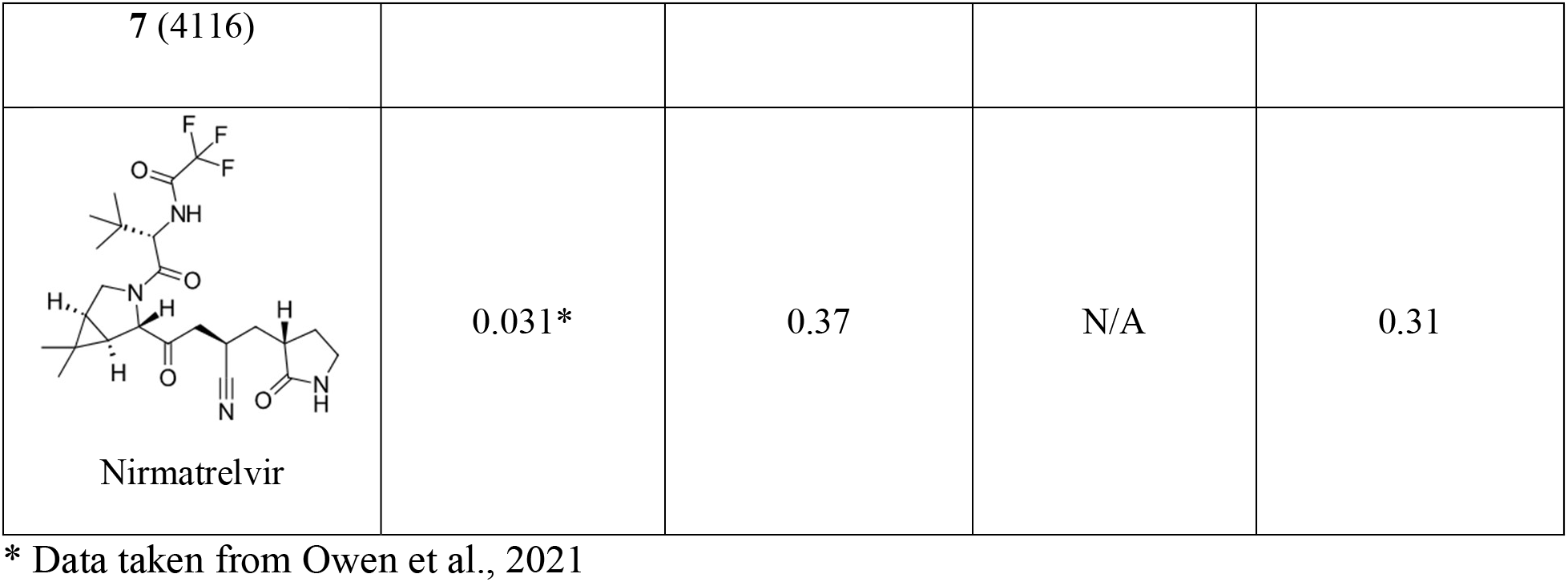
Experimental and predicted *in-silico* inhibition constants (*K*_i_ values) for phenothiazine compounds **1**-**7** and nirmatrelvir with SARS-CoV-2 main protease (M^pro^; PDB: 7L10) [Zhang et al., 2021a,b] and papain-like protease (PL^pro^; PDB: 7CJM) [Fu et al., 2021]. *In-silico K*_i_ values were generated using Molecular Operating Environment 2022.02 (https://chemcomp.com).

| Compounds | SARS-CoV-2 M<sup>pro</sup> |  | SARS-CoV-2 PL<sup>pro</sup> |  |
| --- | --- | --- | --- | --- |
| | Experimental $K_i$ (μM) | Predicted $K_i$ (μM) | Experimental $K_i$ (μM) | Predicted $K_i$ (μM) |
| **1 (4078)** | No Inhibition | 14.53 | No inhibition | 1.59 |
| **2 (4079)** | No Inhibition | 7.71 | No inhibition | 1.61 |
| **3 (4080)** | 80 | 12.86 | No inhibition | 2.39 |
| **4 (4081)** | No Inhibition | 12.92 | No inhibition | 2.26 |
| **5 (4085)** | 10 | 11.09 | 93 | 2.17 |
| **6 (4062)** | 64 | 10.67 | No inhibition | 9.81 |
|  | 51 | 11.47 | No inhibition | 11.74 |
| 7 (4116) |  |  |  |  |
| Nirmatrelvir | 0.031* | 0.37 | N/A | 0.31 |

\* Data taken from Owen et al., 2021

### 2.4 Comparison of M^pro^ and PL^pro^ enzymes of SARS-CoV-2, HCoV-OC43, and HCoV-299E

PDB crystal structures of M^pro^ and PL^pro^ from SARS-CoV-2, OC43 and 229E were required to conduct comparative alignment studies. Crystallization data for M^pro^ of 229E was obtained from the Protein Data bank (PDB: 2ZU2, 1.80 Å) [Lee et al., 2009]; however, crystal structures of OC43 M^pro^ and PL^pro^, and 229E PL^pro^ have not been published. AlphaFold (https://deepmind.com) was therefore utilized to generate predicted structures of the enzymes [Jumper et al., 2021; Varadi et al., 2021]. Primary amino acid sequences corresponding to OC43 M^pro^ (NSP5; AAs 3247-3549), OC43 PL^pro^ (NSP3; AAs 852-2750), and 229E PL^pro^ (NSP3; AAs 898-2484) were obtained from their respective UniProt (https://www.uniprot.org) entries for their coronavirus replicase polyproteins (E.C. 3.4.19.12; E.C. 3.4.22.69; and E.C. 3.4.19.12 respectively). Due to UniProt entries containing the entire NSP3 sequences and not the region that only corresponds to PL^pro^, sequences were aligned with that of SARS-CoV-2 using Clustal Omega (https://www.ebi.ac.uk/Tools/msa/clustalo). Residues that corresponded to the PL^pro^ regions of each NSP (∼300 AAs each) were taken for subsequent generation of AlphaFold structures.

Using the open-access AlphaFold Google Colab notebook (https://colab.research.google.com/github/deepmind/alphafold/blob/main/notebooks/Al phaFold.ipynb) containing a simplified version of AlphaFold 2.3.2, the protein structures of OC43 M^pro^ and PL^pro^ and 229E PL^pro^ was generated from the AA sequence with high model confidence (Supplementary Materials).

Within MOE, the enzyme structures for each CoV strain were individually prepared (Supplementary Materials) and opened within the same window for each protease. Using the *Sequence Editor* window, the *Align/Superpose* feature was utilized. All molecules and atoms not part of the primary sequence (i.e. waters, co-crystallized ligands, etc.) were removed. *Sequence and Structural* alignment was selected with default settings and an alignment executed. Next, superposition of protein structures was competed with default settings (*All Residues* and *Use Current Alignment* selected). The resulting superposition reports containing RMSD values for alignment of all three proteins for each protease can be found in the Supplementary Materials.

### 2.5 Cytotoxicity Assays

Cells seeded into 96-well tissue culture plates were treated with compounds diluted at indicated concentrations and incubated with the cells for 20 hrs. At 20 hrs post-treatment, 10% alamarBlue Cell Viability Reagent (ThermoFisher Scientific, DAL1025) diluted in DMEM/Pen/Strep/Gln containing 2.5% FBS (Huh-7.5 or 293T) was added and further incubated for 4 hrs. Plates were read on FLUOstar Omega 96 well plate reader (BMG Labtech Inc., Ortenberg, Germany) at an excitation of 544 nm and an emission of 580-590 nm. Compound treatments were then normalized to DMSO. To determine the viability of test compounds in Calu-3 cells, a MTT cell viability assay was performed. Calu-3 cells, grown in 96-well plates were treated with two-fold dilutions of the test compound and incubated for 48 hrs. At 48 hrs post-treatment, CellTiter 96® AQueous One Solution Reagent (Promega, G3580), at one-fifth of the culture volume was added to the wells and incubated for 2 hrs. Plates were read at an absorbance of 490 nm with an xMark™ microplate absorbance spectrophotometer (Bio-Rad Laboratories, Hercules, California US). GraphPad PRISM v.10 (https://graphpad.com) was used to determine CC_50_ values by fitting a non-linear curve to the data. The selectivity index (SI) was calculated by dividing the EC_50_ (calculated as described below) by the CC_50_.

### 2.6 HCoV Infection and Quantitation

Viral stocks were prepared as previously described [Pringle et al., 2022]. To test the effect of each phenothiazine derivative (**1**-**7**) on HCoV replication, 293T, Huh-7.5, or Calu-3 cells were infected with HCoV diluted in serum-free Dulbecco’s Modified Eagle Medium (DMEM) at their indicated multiplicity of infection (MOI) at 37 °C for 1 hr. SARS-CoV-2 infected Calu-3 (ATCC HTB-55) cells were plated in DMEM with 10% fetal bovine serum (FBS) and 1% Pen/Strep for two days prior to incubation with an MOI of 0.1 (SARS-CoV-2/Canada/ON/VIDO-01/2020/P2. The inoculum was then removed and replaced with DMEM/Pen/Strep/Gln containing 2.5% FBS for Huh-7.5 and 293T cells, or 2% FBS with 1% Pen/Strep (Calu-3 cells). Cells were supplemented with DMSO, or each phenothiazine derivative at their indicated concentration and maintained for an additional 24 hrs for Huh-7.5 and 293T cells, or for 48 hrs with Calu-3 cells. Viral supernatant was then harvested and stored at -80 °C until titering by median tissue infectious dose (TCID_50_) assay on BHK, Huh7.5, or Vero’76 (ATCC-CRL-1586) cells. SARS-CoV-2 infected cell culture plates were monitored for 5 days for cytopathic effect (CPE). TDC_50_ values were calculated using the Spearman-Kärber method [Hamilton et al., 1977] as previously described [Pringle et al., 2022]. EC50 values and titer plots were generated using PRISM v.10 (https://graphpad.com).

To measure viral genome replication, total RNA was harvested from cells using RNeasy Plus Mini Kit (Qiagen, Hilden, Germany) following the manufacturers’ directions. qPCR was performed using GoTaq 1-Step RT-qPCR (Promega, A6020) reaction, according to the manufacturer’s directions. Primer sequences utilized were as follows: HCoV-OC43 open reading frame (ORF)1a (5’-ATATGGCCAAGGCTGGTGAC-3’ and 5’-ATGTAACACGCCTTCCAGCA-3’), HCoV-229E ORF1a (5’-TGGGTATTGGCGGTCCTAGA-3’ and 5’-GAGCAGTTTCAGGGTCGTCA-3’), and 18S (5’-TTCGAACGTCTGCCCTATCAA-3’ and 5’-GATGTGGTAGCCGTTTCTCAGG-3’).

To visualize replication centers, 293T cells were seeded on #1.5 coverslips (Zeiss, 411025-0003-000) coated with poly-L-lysine (Sigma Aldrich, P2658) and infected the following day with HCoV-OC43 at an MOI of 1. Six hrs after infection, cells were fixed with 4% paraformaldehyde in phosphate buffered saline (PBS) for 15 mins at room temperature. Coverslips were blocked and permeabilized in staining buffer [1% human serum (Sigma Aldrich, 4552) heat-inactivated at 56 °C for 1 hr, 0.1% Triton X-100 in PBS] for 1 hr at room temperature, then stained with J2 anti-dsRNA antibody (Scicons, RNT-SCI-10010200) overnight at 1:500 dilution and 4 °C. The following day the coverslips were washed three times with PBS, then stained with goat anti-mouse Alexa 555 (ThermoFisher, A-21422) in staining buffer for 1 hr at room temperature in the dark. Coverslips were then counterstained for 5 min with Hoescht 33342 (ThermoFisher, 62249), washed three times with PBS, then mounted on coverglass using ProLong Gold anti-fade reagent (ThermoFisher, P36930). Z-stacks were imaged on a Zeiss LSM880 and processed into maximum intensity projections using Zen Black (Zeiss, Oberkochen, Baden-Württemberg, Germany).

## 3.0 RESULTS

### 3.1 Phenothiazines inhibit both SARS-CoV-2 M^pro^ and PL^pro^ activity *in-vitro*

Seven phenothiazine derivatives were evaluated for possible interaction with SARS-CoV-2 M^pro^ (7L10) and PL^pro^ (7CJM). Derivatives **1-6** were synthesized as previously described [Darvesh et al., 2010]. Compound **7** was produced in a good yield (63%) (Supplementary Materials). Spectroscopic methods confirmed molecular structure and purity was confirmed to be >99%.

Enzyme kinetic studies of the phenothiazine derivatives with M^pro^ and PL^pro^ were performed using their respective fluorogenic substrates. Four of the seven compounds (**3**, **5**, **6**, and **7**) showed inhibition of M^pro^ (Table 1). Dose response curves for the phenothiazine derivatives showed that compound **5** exhibited robust inhibition of M^pro^ (Figure 2A). Morrison plots were generated to give *K*_i_ values (Table 1, Figure 2C). Compound **5** was the most potent inhibitor (Figure 2) followed by **7**, **6**, and **3** (Supplementary Materials), while compounds **1**, **2**, and **4** did not inhibit M^pro^ activity. Enzyme kinetic studies with SARS-CoV-2 PL^pro^ showed that only compound **5** inhibited enzyme activity, with a *K*_i_ of 30 µM (Table 1). Representative dose response curves and Morrison plots for compound **5**, are shown in Figure 2B and D respectively. Lineweaver-Burke plots were generated to determine the mode of inhibition for compound **5** with each enzyme (Figure 2E, F). It was shown that compound **5** acts as a competitive inhibitor for M^pro^ and a mixed, non-competitive inhibitor for PL^pro^.

**Figure 2:**
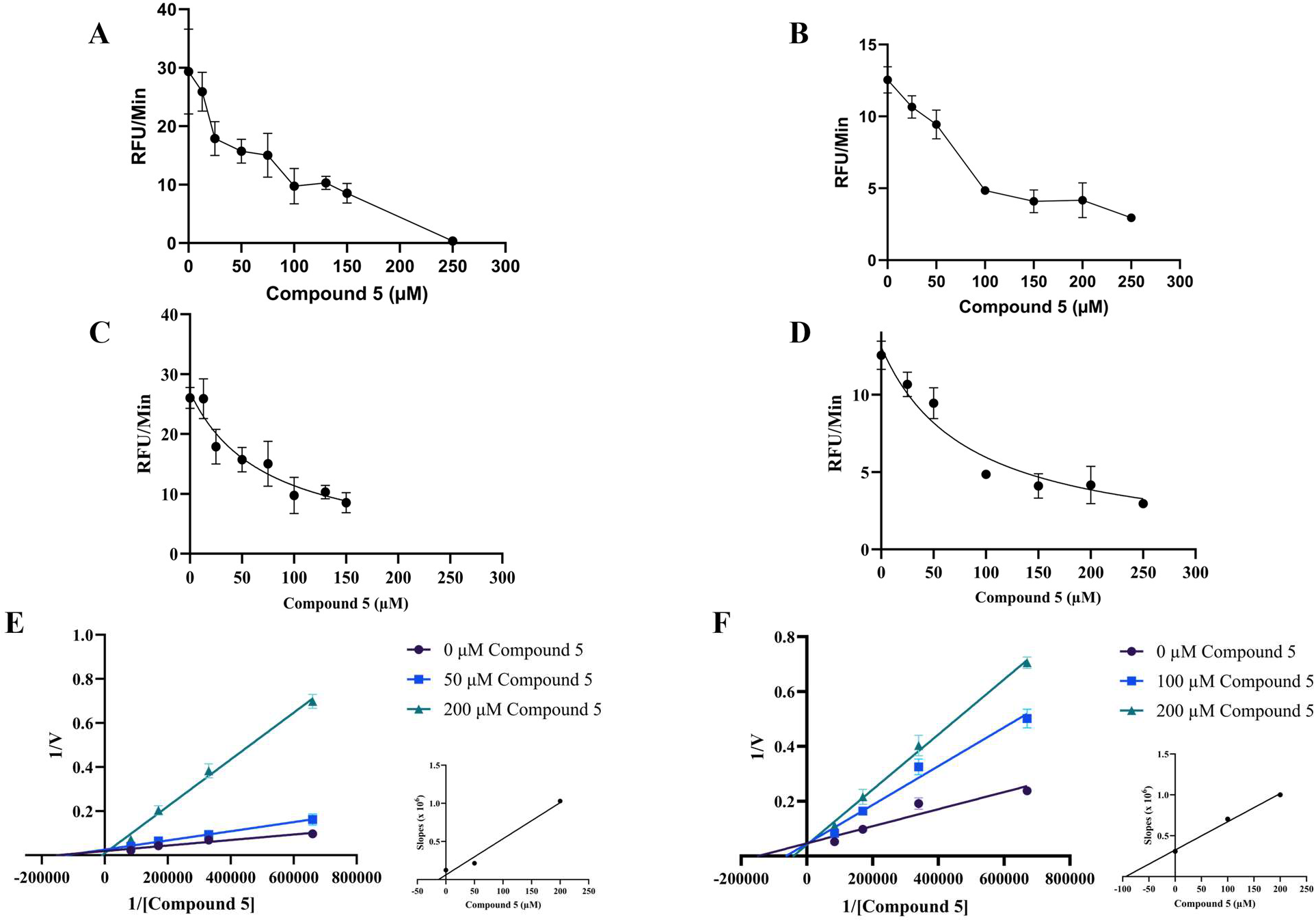
Compound **5** inhibits SARS-CoV-2 main protease (M^pro^) and papain-like protease (PL^pro^). Phenothiazine compound **5** dose response curves, Morrison plots, and Lineweaver-Burk plots for the inhibition of SARS-CoV-2 M^pro^ (**A, C, E** respectively) and SARS-CoV-2 PL^pro^ (**B**, **D**, **F** respectively). Plots were generated using PRISM 9.4.0 (637) (https://graphpad.com).

### 3.2 Binding of phenothiazines to SARS-CoV-2 M^pro^ and PL^pro^

Using a blind docking approach [Hetényi and van der Spoel, 2002; Hetényi and van der Spoel, 2006], all compounds showed propensity to bind to the active site of M^pro^ and desire to bind to the BL2 loop of PL^pro^. Site-specific docking experiments were used to identify the preferred pose (top dock) and interactions within the active site. The predicted *K*_i_s of the preferred pose of all phenothiazine compounds with M^pro^ and PL^pro^ are listed in Table 1. The top docks for each compound with M^pro^ and PL^pro^ were visualized in MOE. Molecular docking data for the most potent compound, **5**, is shown herein, while other compound docking data can be found in the Supplementary Materials.

Overall, docking poses for the phenothiazine derivatives with M^pro^ showed interactions with similar key amino acids (H41, D142, G143, C145, M165, E166, and Q189), many of which were likewise observed in the top docking pose for nirmatrelvir (Supplementary Materials). The compounds that inhibited M^pro^ in experimental assays, **5**, **7**, **6**, and **3,** all showed interactions with at least one of the key amino acids (H41, G143, C145, E166, and Q189) implicated in M^pro^ enzymatic activity [Ngo et al., 2021] (Figure 3A; Supplementary Materials). The most potent phenothiazine, compound **5**, was observed in a self-stabilizing alignment, with the tricyclic phenothiazine moiety folded over itself and forming π-hydrogen bonds (∼4 Å) with the hydrogens surrounding its tertiary amine (Figure 3A). The carbonyl oxygen of compound **5** acts as a hydrogen acceptor, forming bonds with both the SH of the catalytic C145 (3.49 Å) and the NH of G143 (3.06 Å) where it is positioned between them (Figure 3A). Details on docking results of other phenothiazines can be found in the Supplementary Materials.

**Figure 3.**
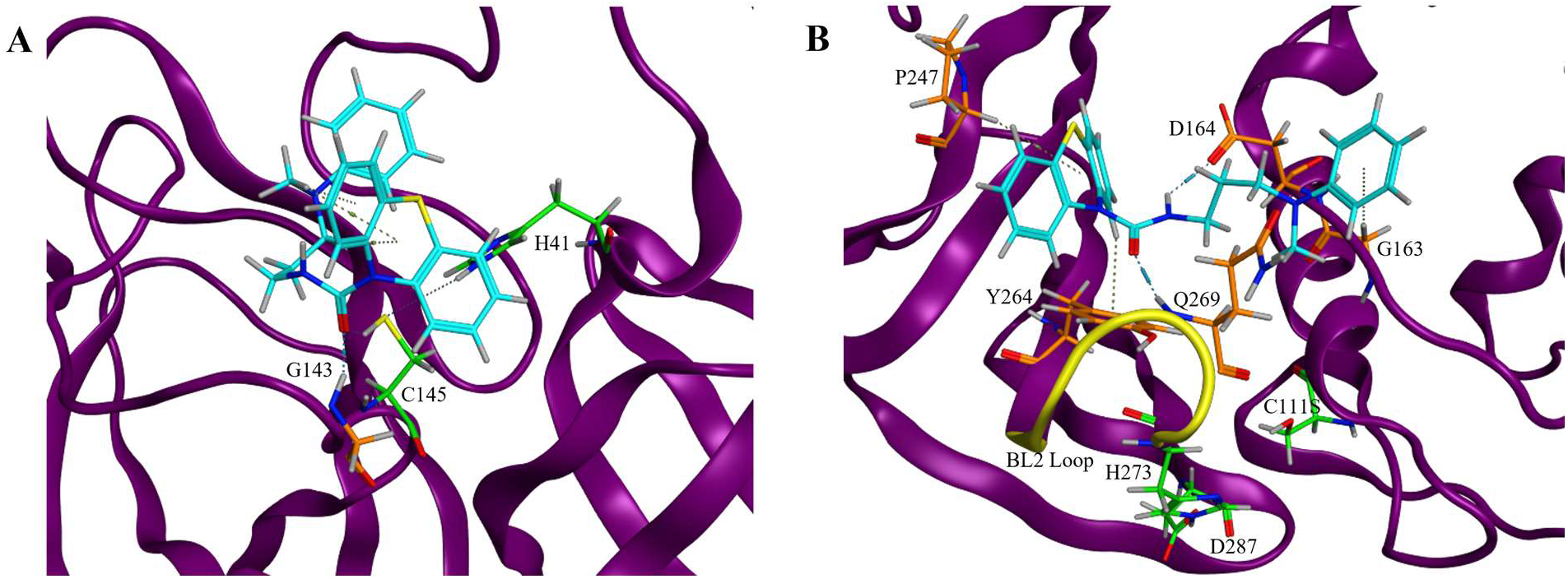
Compound **5** binds to important regions of SARS-CoV-2 main protease (M^pro^) and papain-like protease (PL^pro^) *in-silico*. Top docking poses for compound **5** with SARS-CoV-2 M^pro^ (**A**) and PL^pro^ (**B**). Catalytic amino acids are shown with green carbon atoms (M^pro^: H41 and C145; PL^pro^: C111S, H273, D287), key bonding amino acids are shown with orange carbon atoms (M^pro^: G143; PL^pro^: G163, D164, P247, Y264, Q269), and the SARS-CoV-2 PL^pro^ BL2-loop (AAs 266-272) is shown as the yellow ribbon structure. Images were generated using Molecular Operating Environment 2022.02 (https://www.chemcomp.com).

Only compound **5** demonstrated experimental inhibition of PL^pro^. In its top docking pose, the carbonyl group of compound **5** acts as a proton donor to the NH group of Q269 (3.12 Å) in the BL2 loop (Figure 3B). The phenothiazine moiety points toward the finger domain of PL^pro^ and its center ring forms a π-hydrogen bond with P247 (4.81 Å). An aromatic hydrogen of one of the phenothiazine rings likewise hydrogen bonds with the benzene of Y264 (4.10 Å). The terminal aromatic ring of **5** creates another π-hydrogen bond with the backbone of G163 (4.64 Å). Finally, the secondary amine acts as a proton acceptor with the carbonyl of D164 (2.91 Å).

### 3.3 Homology of M^pro^ and PL^pro^ enzymes of SARS-CoV-2, HCoV-OC43, and HCoV-229E

Sequence and structural alignment of M^pro^ from SARS-CoV-2 (7L10), 229E (2ZU2), and OC43 (AlphaFold PDB) revealed high sequence and structural similarity (Figure 4A) with a total average RMSD value of 1.593 Å, representing the average spatial difference in each AA residue between the three M^pro^ proteins. Furthermore, catalytic amino acids H41 and C145, and key amino acid E166 are highly conserved among these enzymes. Similarly, other key amino acids in the active site of M^pro^, G143 are conserved among all three coronaviruses, while Q189, a key amino acid identified in M^pro^ activity [Ngo et al., 2021], is present in both betacoronaviruses (SARS-CoV-2 and OC43) but replaced by proline in the alphacoronavirus 229E. Detailed alignment reports can be found in the Supplementary Materials.

**Figure 4:**
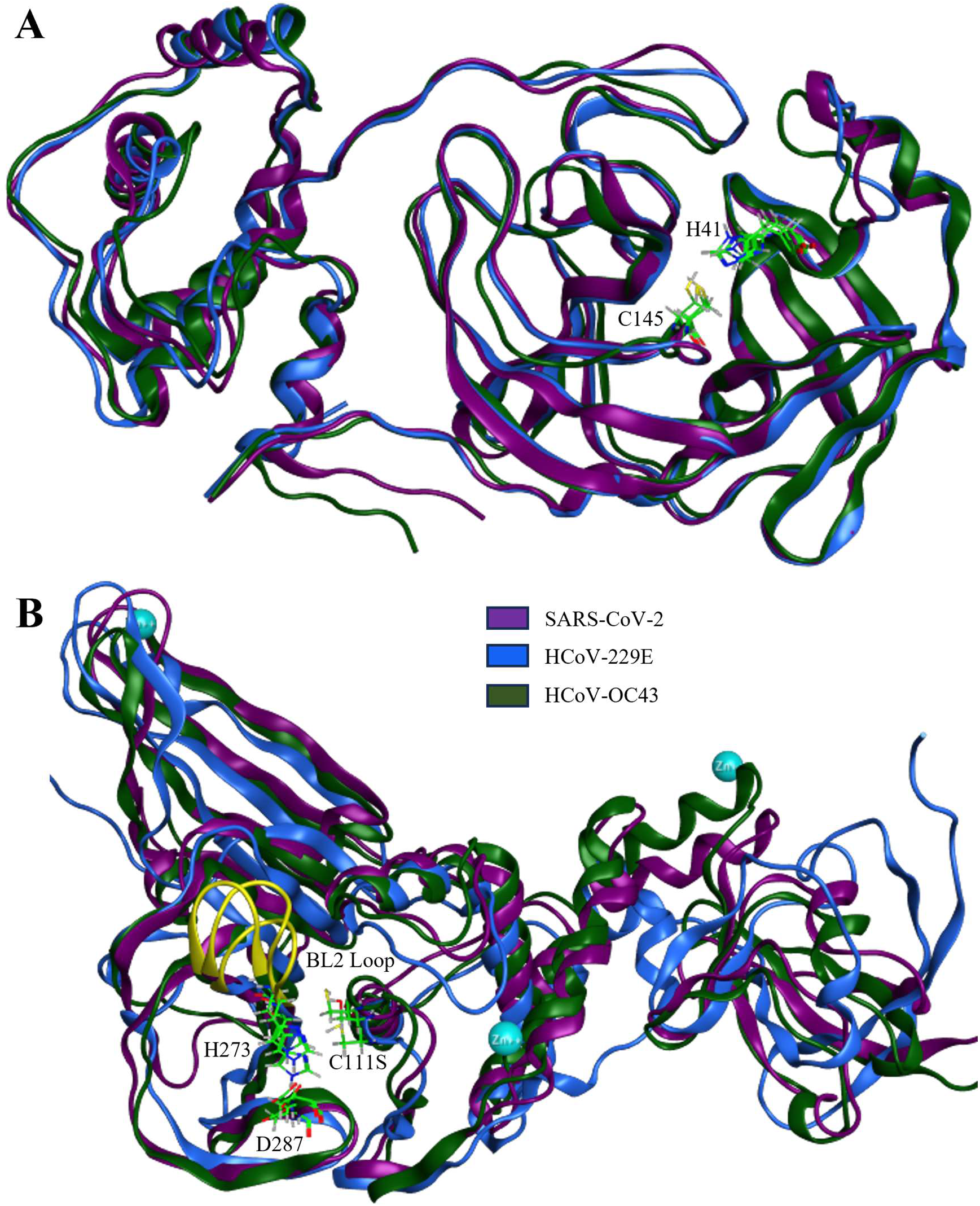
Structural homology shows conservation of cysteine proteases among coronaviruses. Structural alignments of main protease (**A**) and papain-like protease (**B**) among SARS-CoV-2 (purple; PDBs 7L10 [Zhang et al., 2020a.b] and 7CJM [Fu et al., 2021]), HCoV-229E (blue; PDBs 2ZU2 [Lee et al., 2009] and AlphaFold PDB), and HCoV-OC43 (dark green; AlphaFold PDBs). Alignments and images were made using Molecular Operating Environment 2022.02 (https://www.chemcomp.com).

Sequence alignments for PL^pro^ enzymes from SARS-CoV-2 (7CJM) and HCoVs (both AlphaFold PDBs) likewise showed notable sequence homology and conservation of catalytic amino acids C111, H273, and D287 (Figure 4B), with a total average RMSD value of 5.957 Å. Amino acids corresponding to the BL2 loop of SARS-CoV-2 PL^pro^ (AAs 266-272) showed partial homology among all three coronaviruses, with the two glycine residues (G267 and G272) being conserved. Detailed alignment reports can be found in the Supplementary Materials.

### 3.5 Compound 5 inhibits replication of OC43, 229E, and SARS-CoV-2 *in-cellulo*

Seven phenothiazine compounds were tested in an OC43 infection model. Consistent with enzyme kinetic studies and *in-silico* analyses, compound **5** showed the greatest degree of inhibition (Figure 5A; EC_50_ = 8.95 µM, SI >40). Compound **5** was further tested in 229E and SARS-CoV-2 infection models and induced a similar degree of inhibition, but with less cytotoxicity at inhibitory concentrations (Figure 5B, C; EC_50_ = 3.31 µM, SI = 15.9; EC_50_ = 1.08 µM, SI >40 respectively).

**Figure 5.**
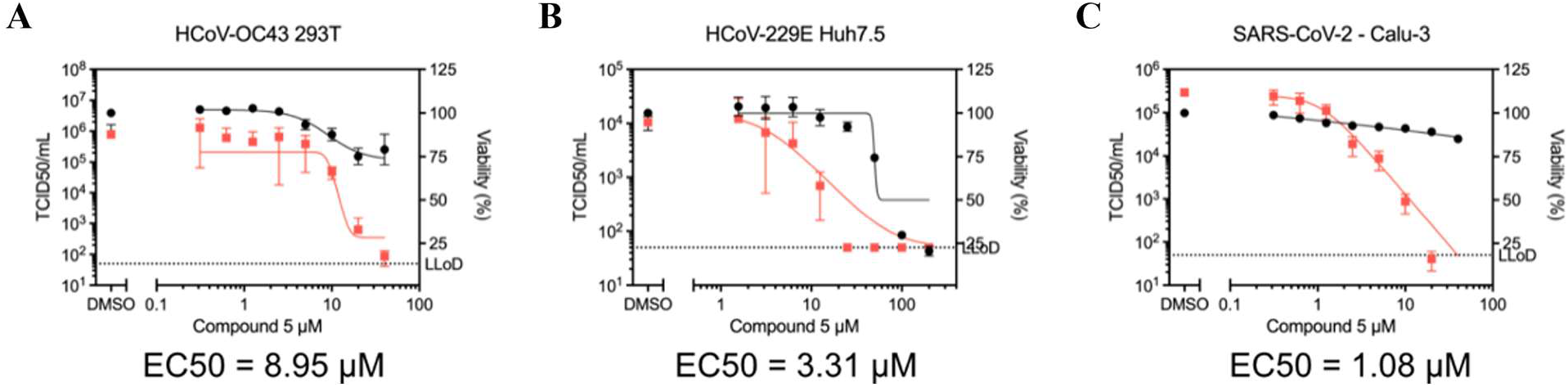
Compound **5** reduces viral titer in several coronavirus models. Phenothiazine compound **5** tested in HCoV-OC43 (**A**), HCoV-229E (**B**), and SARS-CoV-2 (**C**) infection models. Viral titer was determined by 50% tissue culture infectious dose (TCID_50_) assay and is shown as red squares. HEK293T cells were infected with HCoV-OC43 at an MOI of 0.1 then treated with **5** for 24 hrs before harvest and assayed with BHK cells. Huh7.5 cells were infected with HCoV-229E at an MOI of 0.1 then treated with **5** for 24 hrs before harvest and assayed with Huh7.5 cells. Calu-3 cells were infected with SARS-CoV-2 at an MOI of 0.1 then treated with **5** for 48 hrs before harvest and assayed using Vero’76 cells. Percent cell viability of uninfected cells is shown as black circles and was measured in parallel. All experiments were completed in triplicate and their average values shown with standard error bars. EC_50_ values and plots were generated using PRISM 9.4.0 (637) (https://graphpad.com).

Inhibition of viral proteases should inhibit early steps in infection and inhibit the formation of cytoplasmic replication organelles (ROs). To test this, we infected cells with OC43 at a high MOI then added compound **5** at one hour after infection. At 6 hrs post-infection, numerous ROs were found in the cytoplasm by staining with the J2 monoclonal antibody for dsRNA, a necessary replication intermediate of RNA viruses. Addition of compound **5** at inhibitory concentrations led to a loss of dsRNA staining (Figure 6A). Concordantly, full-length viral genomes fail to accumulate in the presence of compound **5** in both OC43 and 229E infection models (Figure 6B, C).

**Figure 6.**
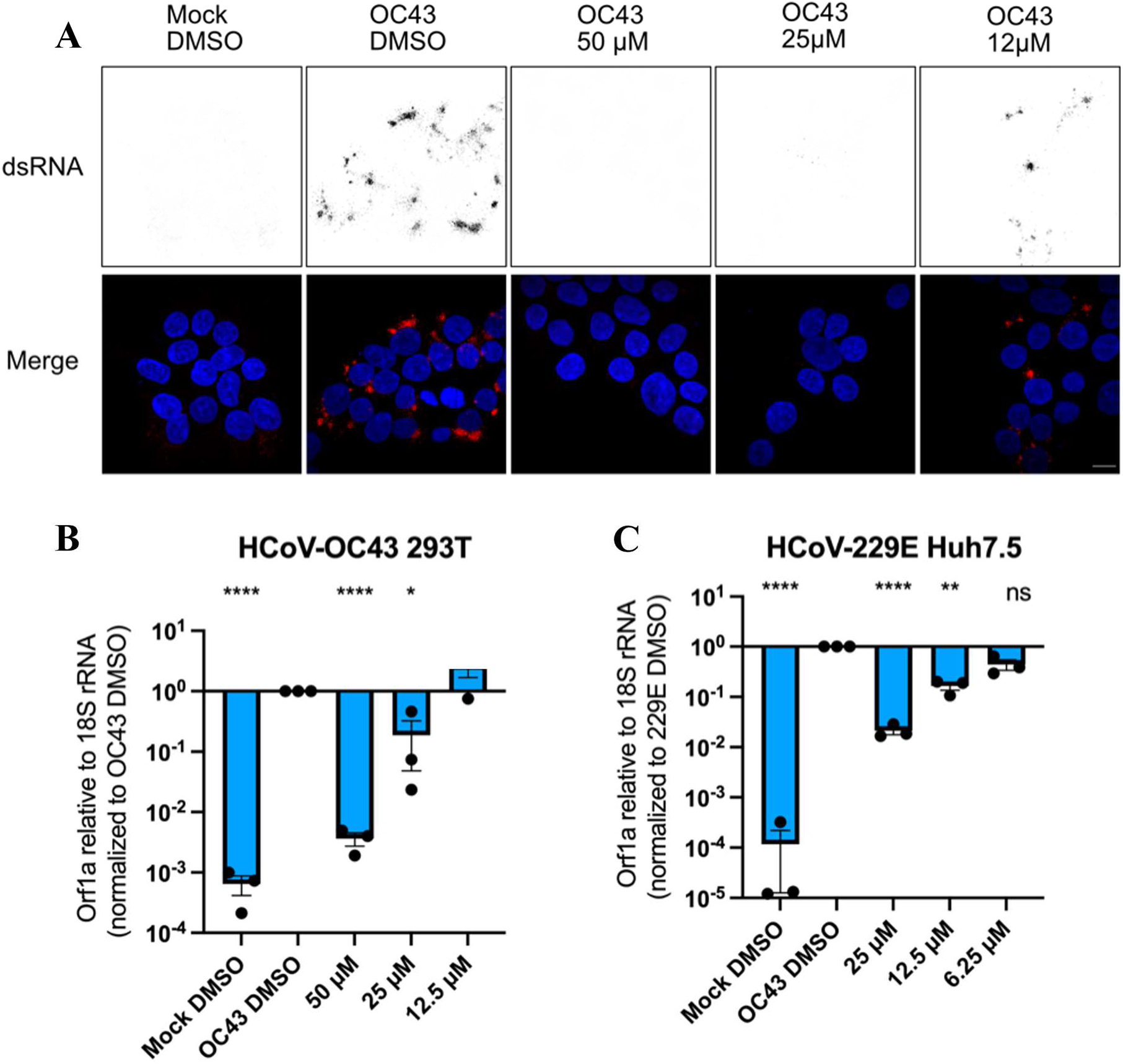
Compound **5** inhibits viral RNA production. Visualization of the formation of viral replication machinery when treated with phenothiazine compound **5** at varying concentrations in HCoV-OC43 infected cells (**A**). HEK293T cells were seeded on coverglass and infected with HCoV-OC43 at an MOI of 1, then treated with compound **5** or vehicle control for 6 hrs before harvest. Cells were fixed and stained with a monoclonal antibody that detects dsRNA. Scale bar = 10 μm. Quantification of formation of viral genetic material when treated with **5** in HCoV-OC43 (**B**) and HCoV-229E (**C**) infection models. HEK293T cells were infected with HCoV-OC43 at an MOI of 0.1 then treated with compound **5** for 24 hrs before harvest. Huh7.5 cells were infected with HCoV-229E at an MOI of 0.2 then treated with compound **5** for 24 hrs before harvesting. Viral genomes were quantified by RT-qPCR in triplicate and average values shown with standard error bars. **** = p<0.0001; ** = p<0.01; * = p <0.05; ns, non-significant by Students’ t-test compared to infected DMSO-treated.

## 4.0 DISCUSSION

Despite the protection afforded by SARS-CoV-2 vaccines, antivirals still play a critical role in treatment of immune escape variants [Zolfaghari et al., 2022; Iketani et al., 2023]. Furthermore, vaccines are not yet available for other known HCoVs or pre-emergent coronaviruses with pandemic potential. Currently, there are two clinically approved antiviral drugs widely used for the management of COVID-19 symptoms in mild-to-severe cases, Paxlovid™ (nirmatrelvir and ritonavir) and remdesivir [Reis et al., 2022; Eastman et al., 2020; Grundeis et al., 2023]. Despite their approval, double-blind and placebo-controlled studies with expansive and diverse subject groups remain limited, and many intervention methods were approved for use under emergency need; thus, their long-term effects and efficacy are not yet well-understood [Reis et al., 2022; Grundeis et al., 2023]. Nirmatrelvir irreversibly inhibits SARS-CoV-2 M^pro^ by forming a covalent bond with its catalytic cysteine [Li et al., 2022]. As this drug is rapidly metabolized by cytochrome P450, it is co-formulated with ritonavir, a cytochrome P450 3A4 (CYP3A4) inhibitor, to make the drug cocktail, Paxlovid™, which is currently the front-line antiviral treatment for SARS-CoV-2 infection [Hammond et al., 2022; Food and Drug Administration, 2022; Owen et al., 2021]. Furthermore, both Paxlovid™ and remdesivir have been shown to cause severe adverse effects in some patients [Hammond et al., 2022; Reis et al., 2022; Tian et al., 2023], particularly in patients with medicated comorbidities who are at higher risk for COVID-19 complications [Cao et al., 2023; Prikis and Cameron, 2022].

Herein, phenothiazine compounds **1**-**7** were assessed for their potential to inhibit multiple SARS-CoV-2 enzymes. The proteolytic activity of both SARS-CoV-2 M^pro^ and PL^pro^ enzymes were inhibited *in-vitro* by compound **5**, as was *in-cellulo* viral replication in SARS-CoV-2, OC43, and 229E infection models. Some other compounds inhibited proteolysis of M^pro^ to a lesser degree, but only compound **5** showed the unique ability to inhibit both enzymes. Currently, it is unclear whether the observed *in-vivo* inhibitory effects of compound **5** are a result of its activity against M^pro^, PL^pro^, or both.

*In-silico* assessment of the phenothiazines with M^pro^ highlight key AAs previously identified [Zhang et al., 2020a,b; Ngo et al., 2021]. Most notably, compound **5** is the most potent inhibitor, likely due to its binding interactions with C145 and G143 (Figure 3), which are also observed in the binding of nirmatrelvir [Li et al., 2022]. This may indicate that these residues are essential targets for antiviral intervention of this enzyme. Docking studies of **5** with PL^pro^ showed interactions with the BL2 loop that were likewise observed in other PL^pro^ inhibitor studies [Fu et al., 2021; Ratia et al., 2008]. Specifically, bonding with Q269 seems to be key here. Further studies are required to understand how the BL2 loop and its residues impact the activity of PL^pro^ and CoVs in general.

Evaluation of the sequential and structural similarities of M^pro^ and PLpro enzymes among SARS-CoV-2, OC43, and 229E (Figure 4A, B) suggests conservation of residues involved in enzymatic activity. M^pro^ showed a higher degree of sequential and structural conservation, notable of key AAs (C145, H41, G143, M165, E166) [Zhang et al., 2020a,b; Ngo et al., 2021], while PL^pro^ showed greater structural and sequential variability, likely due to its additional activities in ubiquitin metabolism and host immune function [Shin et al., 2021], however, it’s catalytic domain and BL2 loop still showed notable conservation (C111, H273, D287, G267, and G272). Other studies have also explored the homology of PL^pro^ among human coronaviruses and those of other organisms showing similar findings [Shin et al., 2021; Ullrich and Nitsche, 2022]. These results suggest that compounds determined to interact with either of these proteases within a specific CoV strain or host organism, may additionally be active against those of other CoVs. This indicates potential for development of pan-coronavirus antivirals that could have applications in both human and veterinary medicine, as well as may aid in the treatment of yet undiscovered coronaviruses.

## 5.0 CONCLUSIONS

These findings suggest that the phenothiazine urea compound, **5** causes potent inhibition of the viral replication of SARS-CoV-2, HCoV-OC43, HCoV-229E, and may broadly inhibit other diverse coronaviruses. Phenothiazine containing drugs should be investigated further for the treatment of coronavirus-mediated disease. Furthermore, due to high conservation of M^pro^ and PL^pro^ enzymes, coronaviruses such as OC43 and 229E can be used as models to rapidly screen antivirals for more virulent coronavirus strains.

## Abbreviations

M^pro^: Main Protease
PL^pro^: papain-like protease
MOE: Molecular Operating Environment
*K*_i_: inhibition constant

## Acknowledgements

This work was supported in part by the Canadian Institutes of Health Research (PJT – 153319; VS1-175531), the Canadian Foundation for Innovation (Grant No. 37854), the Dalhousie Medical Research Foundation (DMRF Chemists; The Durland Breakthrough Fund; DMRF Clare Durland Fund in Alzheimer’s Disease Research; DMRF Research Grant – Robert and Barbara Pickett; DMRF Gillian’s Hope for MS Research Fund), and the Dalhousie Medical Research Foundation Irene MacDonald Sobey Endowed Chair in Curative Approaches to Alzheimer’s Disease. Mitacs Accelerate Grant (IT29621 and IT31142) and partner organization, Applied Pharmaceutical Innovation (API), Edmonton, Alberta. VIDO receives operational funding from the Government of Saskatchewan through Innovation Saskatchewan and the Ministry of Agriculture and from the Canada Foundation for Innovation through the Major Science Initiatives for its CL3 facility.

## Author contributions

*Katrina Forrestall:* Data Curation, Formal Analysis, Investigation, Methodology, Software, Valiation, Visualization, Writing – original draft, Writing – review and editing.

*Eric S. Pringle*: Formal analysis, Investigation, Writing – original draft, Writing – review & editing.

*Dane Sands*: Formal Analysis, Investigation, Methodology, Writing – original draft, Visualization, Writing – review & editing.

*Brett Duguay*: Formal analysis, Investigation

*Brett Farewell*: Investigation, Visualization, Writing – original draft, Writing – review & editing

*Tekeleselassie Woldemariam*: Investigation, Formal Analysis, Visualization, Writing – review & editing

*Darryl Falzarano*: Funding Acquisition, Investigation, Methodology, Resources, Supervision, Writing – review & editing

*Ian Pottie*: Conceptualization, Formal Analysis, Funding Acquisition, Methodology, Project Administration, Resources, Supervision, Visualization, Writing – original draft, Writing – review & editing

*Craig McCormick*: Conceptualization, Formal Analysis, Funding Acquisition, Methodology, Project Administration, Resources, Supervision, Visualization, Writing – original draft, Writing – review & editing

*Sultan Darvesh*: Conceptualization, Formal Analysis, Funding Acquisition, Methodology, Project Administration, Resources, Supervision, Visualization, Writing – original draft, Writing – review & editing

## Data Availability Statement

All data is available from the corresponding author upon reasonable request.

## Notes

### Competing Interest Statement

Sultan Darvesh is a scientific cofounder and stockholder in Treventis Corporation, a biotech company focused on development of diagnostic and therapeutic agents for Alzheimers disease. Sultan Darvesh and Ian R. Pottie are listed as inventors on patents assigned to Treventis Corporation. The other authors declare there are no competing interests.

