## Supplemental Materials for "Dual inhibition of coronavirus M^pro^ and PL^pro^ enzymes by phenothiazines and their antiviral activity"

**TABLE OF CONTENTS**

**S1.0 EXPANDED MATERIALS AND METHODS** .......... **S3**
**S1.1 Phenothiazine Compounds** .......... S3
*S1.1.1 Spectroscopic data of* N-(Adamant-2-yl)-10H-phenothiazine-10-carboxamide .......... S3
**S1.2 *In-silico* Docking Studies** .......... S9
*S1.2.1 Preparation of Enzyme PDB Files for Docking* .......... S9
*S1.2.2 Preparation of Compounds for Docking* .......... S9
*S1.2.3 Docking Procedure* .......... S10
*S.1.2.4 Validation of Docking Procedure* .......... S10
**S1.3 Generation of AlphaFold PDB Structures for HCoV-OC43 and HCoV-229E** .......... S11
*S1.3.1 Generation of $M^{pro}$ structures* .......... S11
*S1.3.2 Generation of $PL^{pro}$ structures* .......... S11
**S2.0 EXPANDED RESULTS** .......... **S13**
**S2.1 Enzyme Kinetics** .......... S13
**S.2.2 *In-silico* Docking Studies** .......... S14
**S2.3 Homology of $M^{pro}$ and $PL^{pro}$ Enzymes of SARS-CoV-2, HCoV-OC43, and HCoV-229E** .......... S16
*S2.3.1 Sequential Alignments* .......... S16
*S2.3.2 Structural Alignments* .......... S17
**S2.4 $TCID_{50}$ Assay** .......... S18
**S3.0 REFERENCES** .......... **S19**

### S1.0 MATERIALS AND METHODS EXPANDED

#### S1.1 Phenothiazine Compounds

Phenothiazine derivative, **7** (Figure S1) was synthesized following a previously used method [Darvesh et al., 2010] but with condensation of phenothiazine-10-carbonyl-chloride (Thermo Fisher Scientific Chemicals; https://www.thermofisher.com) and 2-aminoadamantane hydrochloride (Sigma Aldrich, 153818). Phenothiazine-10-carbonyl chloride (0.3926 g, 1.5 mmol) was added to dichloromethane (15 mL) and cooled in an ice bath. Triethylamine (0.523 mL, 3.75 mmol) was added dropwise, followed by 2-aminoadamantane hydrochloride (0.422 g, 2.25 mmol). The mixture was warmed to room temperature and stirred for 3 hrs. The reaction mixture was washed with $H_2O$ (2 × 20 mL), 0.1 *N* $HCl_{(aq.)}$ (2 × 20 mL), and again with $H_2O$ (2 × 20 mL). The organic layer was dried over $Na_2SO_4$, gravity filtered, and concentrated *in vacuo* to afford *N*-(adamant-2-yl)-*10H*-phenothiazine-10-carboxamide as a white solid (0.354 g, 0.941 mmol, 69% yield).

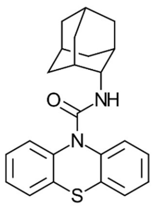


**Figure S1:** *N*-(Adamant-2-yl)-10*H*-phenothiazine-10-carboxamide

*S1.1.1 Spectroscopic data of* N-(Adamant-2-yl)-10H-phenothiazine-10-carboxamide

MP 144.2-147.4 ºC; IR(ATR) 3428, 3066, 2892, 2848, 1682, 1496, 1460, 1248, 1218, 754, 734 $cm^{-1}$; $^1$H NMR (400 MHz, $CDCl_3$) δ 7.60 (dd, J = 8.0, 1.1 Hz, 2H), 7.40 (dd, J = 7.8, 1.4 Hz, 2H), 7.32 (td, J = 7.4, 1.4 Hz, 2H), 7.20 (td, J = 7.6, 1.2 Hz, 2H), 5.34 (td, J = 7.3 Hz, 1H), 3.97-3.95 (m, 1H), 1.98-1.91 (m, 2H), 1.86-1.78 (m, 5H), 1.77-1.72 (m, 1H), 1.72-1.68 (m, 2H), 1.61-1.53 (m, 2H), 1.50-1.43 (m, 2H); $^{13}$C NMR (100.7 MHz, $CDCl_3$) δ 153.9, 139.0, 133.5, 128.2, 127.2, 126.6, 54.9, 37.6, 37.2, 32.2, 27.3, 27.2; HRMS (ESI$^+$): calculated for $C_{23}H_{24}N_2NaOS^+$: 399.1502 amu; found for $C_{23}H_{24}N_2NaOS^+$: 399.1502 amu; HPLC purity at 230 nm ($CH_3CN$ 100%, retention time: 4.956 mins) 100%.

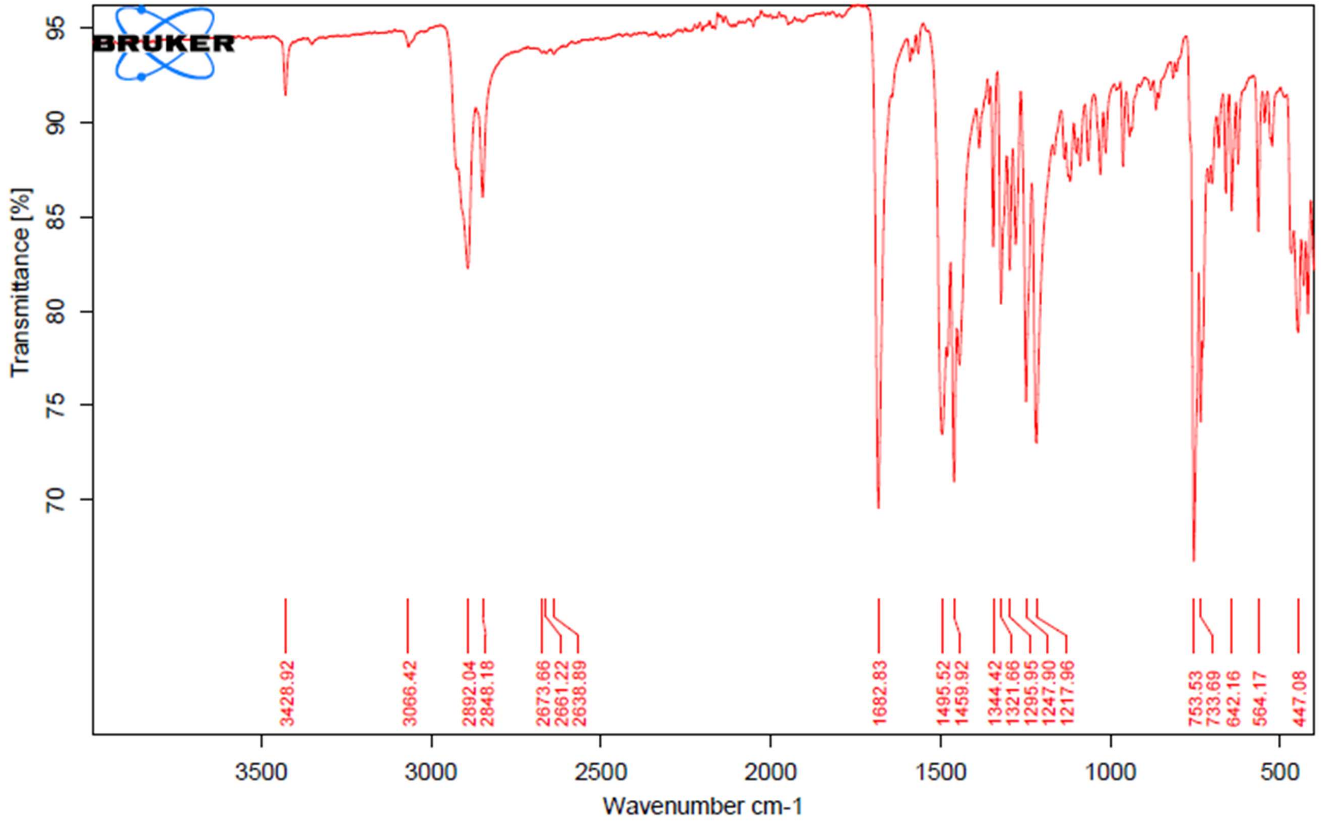


**Figure S2:** *N*-(Adamant-2-yl)-10*H*-phenothiazine-10-carboxamide IR(ATR)

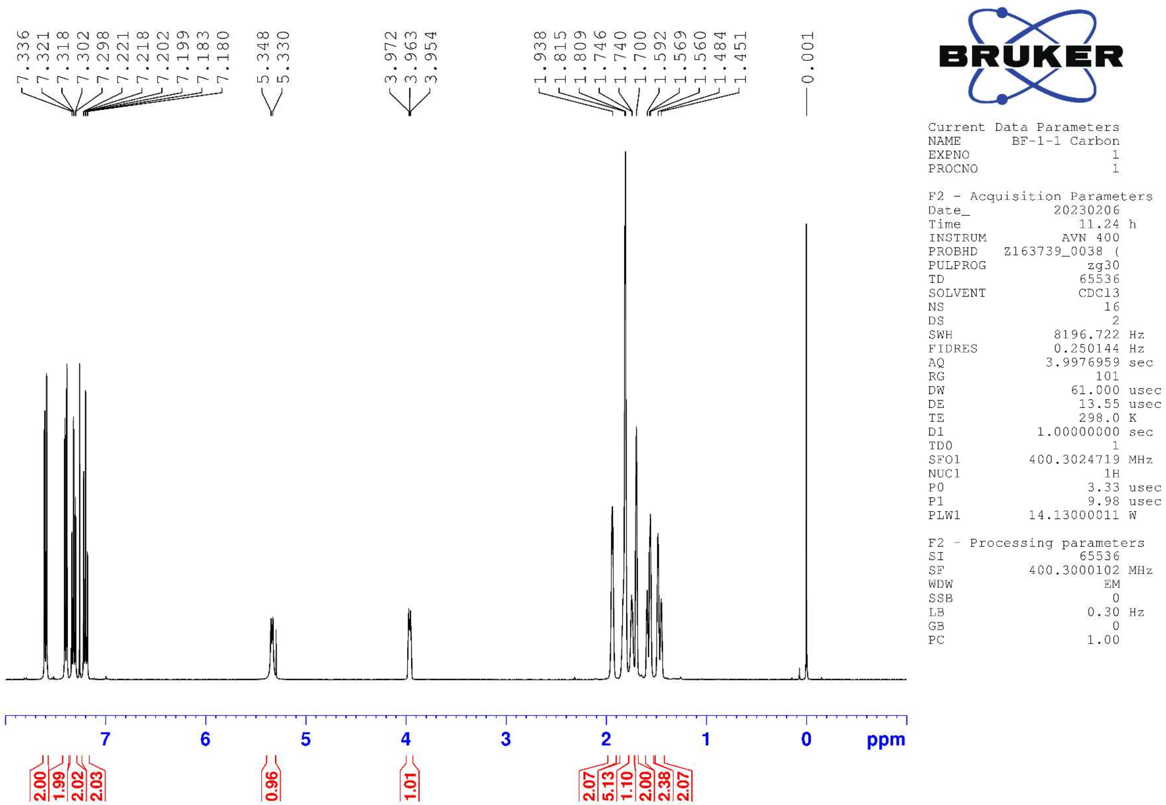


**Figure S3:** *N*-(Adamant-2-yl)-10*H*-phenothiazine-10-carboxamide $^1$H NMR 400 MHz ($CDCl_3$)

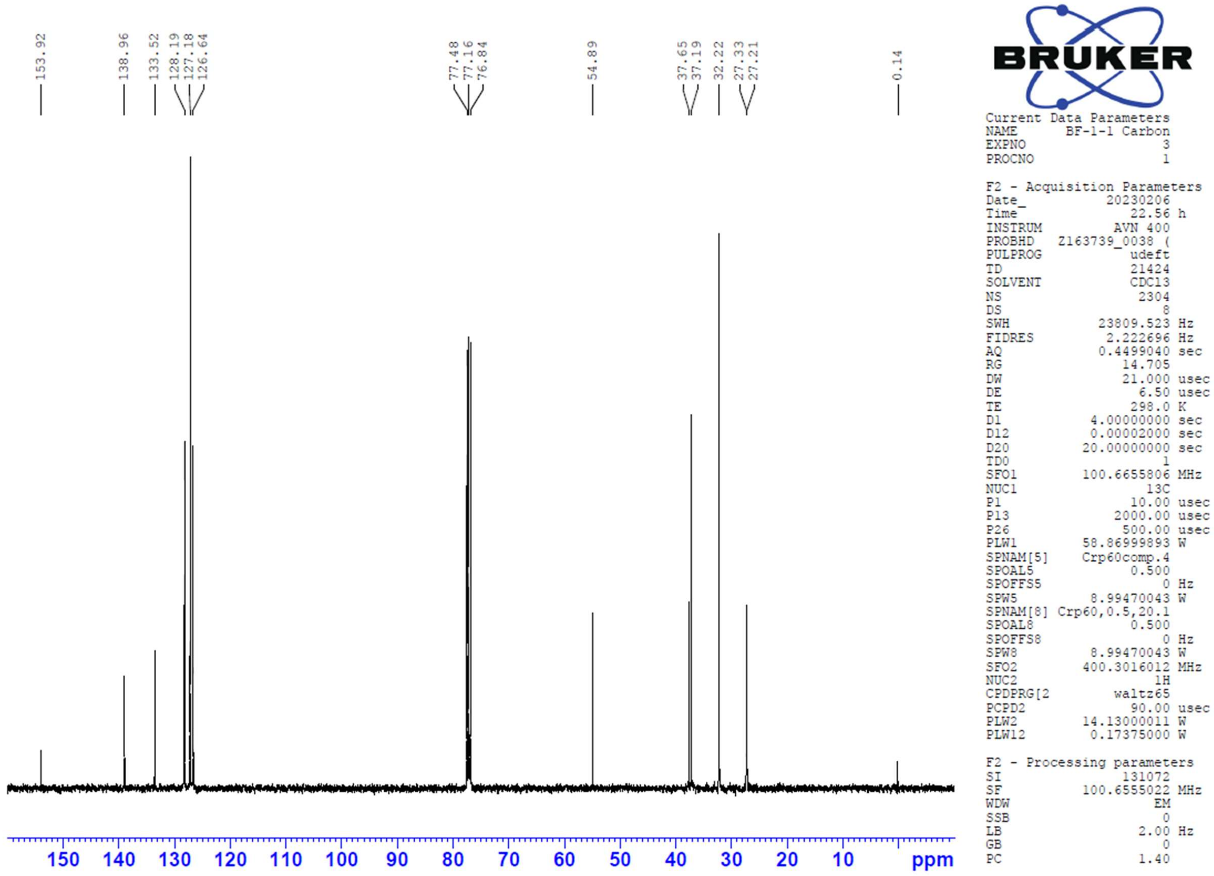


**Figure S4:** *N*-(Adamant-2-yl)-10*H*-phenothiazine-10-carboxamide $^{13}C$ udeft NMR 100.7 MHz ($CDCl_3$)

### Mass Spectrum SmartFormula Report

**Analysis Info**

| | | | |
|---|---|---|---|
| Analysis Name | D:\Data\Xiao\Feb 17 2023000010.d | Acquisition Date | 2023-02-17 9:53:40 AM |
| Method | Xiao all 1.m | Operator | x |
| Sample Name | N-2-Adamantanyl-10-phenothiazinecarboxamide | Instrument | compact 8255754.20059 |
| Comment | | | |

**Acquisition Parameter**

| | | | | | |
|---|---|---|---|---|---|
| Source Type | ESI | Ion Polarity | Positive | Set Nebulizer | 0.5 Bar |
| Focus | Not active | Set Capillary | 3500 V | Set Dry Heater | 180 °C |
| Scan Begin | 50 m/z | Set End Plate Offset | -500 V | Set Dry Gas | 4.0 l/min |
| Scan End | 1500 m/z | Set Charging Voltage | 2000 V | Set Divert Valve | Source |
| | | Set Corona | 0 nA | Set APCI Heater | 0 °C |

| Meas. m/z | Ion Formula | m/z | err [ppm] |
|---|---|---|---|
| 399.1502 | C23H24N2NaOS | 399.1502 | -0.2 |

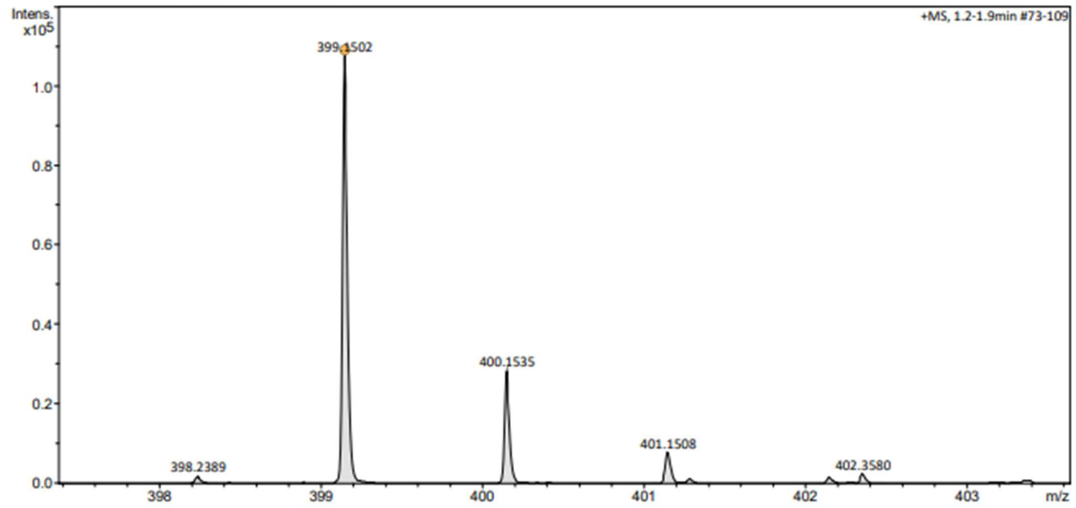


**Figure S5:** *N*-(Adamant-2-yl)-10*H*-phenothiazine-10-carboxamide High Resolution Mass Spectroscopy (ESI, positive mode)

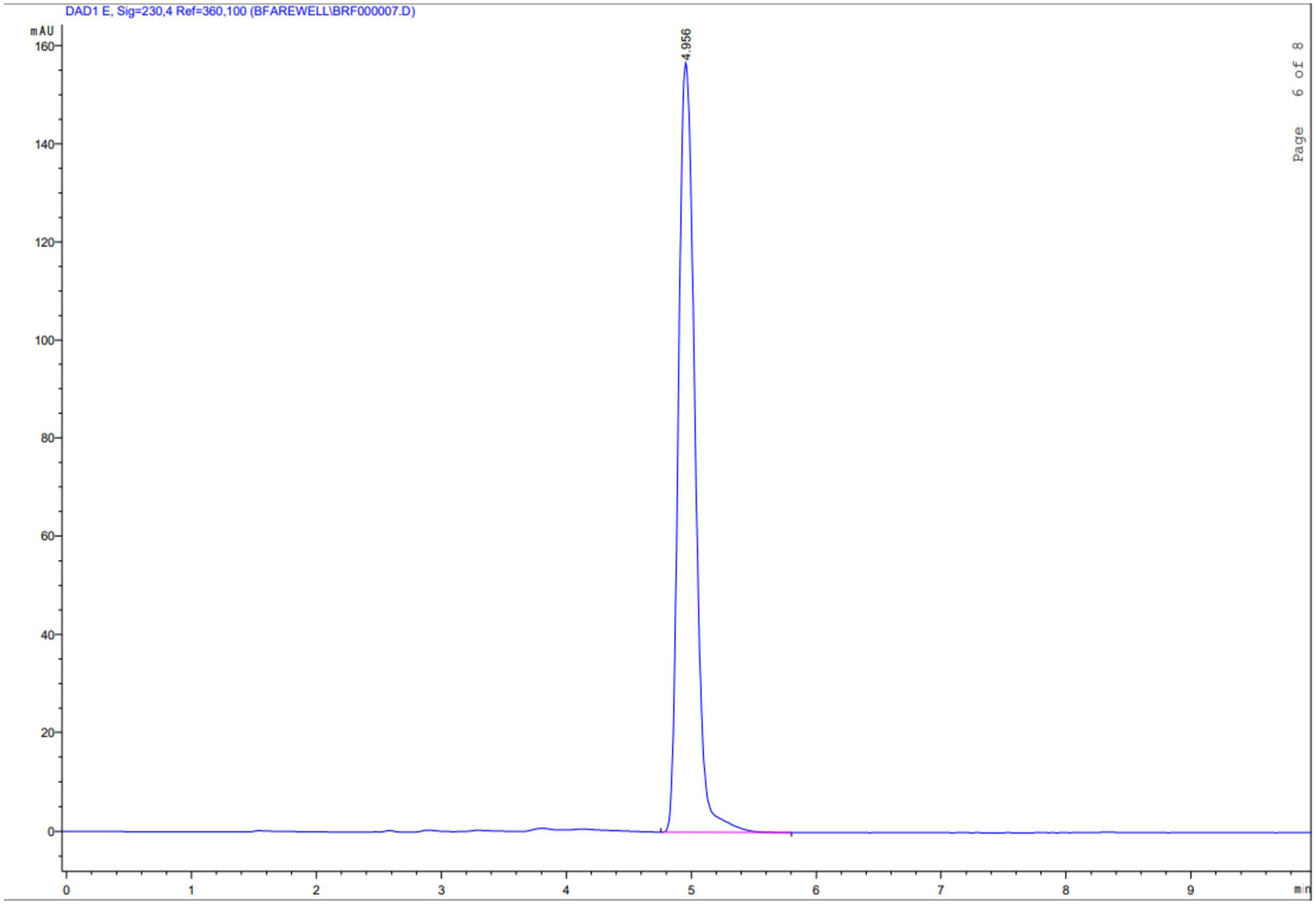


**Figure S6:** *N*-(Adamant-2-yl)-10*H*-phenothiazine-10-carboxamide HPLC Trace

### S1.2 *In-silico* Docking Studies

*S1.2.1 Preparation of Enzyme PDB Files for Docking*

Crystal structures of SARS-CoV-2 $M^{pro}$ (PDB: 7L10, 1.63 Å) [Zhang et al., 2020a,b] and $PL^{pro}$ (PDB: 7CJM, 3.20 Å) [Fu et al., 2021] were obtained from the Protein Databank (https://rcsb.org) and prepared for molecular docking. Within Molecular Operating Environment (MOE), each enzyme monomer was loaded individually and assessed within the *Sequence (Seq)* editor window. All ligands and atoms non-essential to protein function were removed ($Zn^{2+}$ atoms were retained for $PL^{pro}$), leaving the enzyme amino acid sequence and water molecules. Using the MOE *Structure Preparation* feature, corrections were made to the enzyme to repair charge errors, input missing residues, and fix residues with atoms containing fractional occupancies. Next, *Prononate3D* with default settings (pH 7.0; Temperature 300 K; Salt Concentration: 0.1 M; Electrostatics: GB/VI; Dielectric: 2; van der Waals: 800R3) was used to simulate an aqueous environment. Final corrections to system charges were made using the *Structure Preparation* feature once again. Each prepped enzyme was saved as a .moe file and used for subsequent docking.

*S1.2.2 Preparation of Compounds for Docking*

The chemical structure of each compound was individually constructed within MOE using the *Builder* window with DaylightSMILES input. SMILES strings for each compound were generated within ChemDoodle 2D 11.3.0 (https://chemdoodle.com). Once built, each compound was minimized using the MOE *Minimize* feature with the default Amber10 Extended Hückle Theory (Amber10:EHT) forcefield. Compounds were input into a MOE database file and prepared within the database window as follows: *Compute* > *Molecule* > *Wash* with default settings at pH 7.0; *Compute* > *Molecule* > *Energy Minimize* with default settings; and *Compute* > *Molecule* > *Conformational Search* with default settings but with the *Rejection Limit* changed to 500. This generated a new database containing conformers of each compound, which was then used for subsequent docking.

*S1.2.3 Docking Procedure*

In general, docking procedures remained the same between blind and site directed docks, except where *Site* was identified as *Receptor Atoms* or *Dummy Atoms* for blind and site docks respectively. All other docking settings used were: *Receptor*: prepared enzyme file, *Receptor and Solvent Atoms* selected; *PH4*: *None*; *Density*: no file selected; *Ligand*: prepared conformer database, *Ligand Atoms* selected, *Conformers*: *use existing*; *Placement*: *Triangle Matcher* with timeout set to 300 and number of returned poses set to 10000, *Score*: *London dG*, and *Poses*: 300; *Refinement*: *Induced Fit* with default settings, *Score*: *GBVI/WSA dG*, and *Poses*: 5.

To generate dummy atoms for site-directed docking, the MOE *Site Finder* feature was utilized. With a prepped enzyme file loaded in the main MOE window, *Compute > Site Finder > Apply* was selected. Using the generated list of sites in the *Site Finder* viewer, blind docking results were browsed in MOE and their binding locations assigned to its corresponding *Site Finder* sites. If a site was identified as the active site (or in the case of $PL^{pro}$, sites corresponding to the active site and BL2 loop together), *Dummies* was selected within the *Site Finder* window and dummy atoms were generated in the enzyme file. This file was then saved as a prepared enzyme .moe file and used for subsequent site-directed docking as described above.

*S.1.2.4 Validation of Docking Procedure*

Crystal structures for $M^{pro}$ (7L10) and $PL^{pro}$ (7CJM) were selected based on their previous use with *in-silico* studies [Zhang et al. 2020a,b; Fu et al. 2021] and the fact that each PDB contained a co-crystallized ligand molecule, of similar aromatic character to compounds assessed in this study, that could be exploited under a self-docking protocol to generate a Root Mean Square Deviation (RMSD) value for validation of docking. An RMSD value <2.0 Å, representing the angstrom difference in ligand placement between a crystallized ligand and its self-dock is considered appropriate validation of docking procedure [Veith et al., 1998; Bursulaya et al., 2003]. The docking procedure used for self-docking of each enzyme with its co-crystallized ligand was the same as the site-directed docking procedure described above. Both $M^{pro}$ and $PL^{pro}$ enzymes were validated with an RMSD of 0.044 Å and <0.000 Å respectively.

**S1.3 Generation of AlphaFold PDB Structures for HCoV-OC43 and HCoV-229E**

*S1.3.1 Generation of $M^{pro}$ structures*

The open-access AlphaFold Google Colab notebook (https://colab.research.google.com/github/deepmind/alphafold/blob/main/notebooks/AlphaFold.ipynb), containing a simplified version of AlphaFold 2.3.2, was used to generate a predicted crystal structure of OC43 M$^{pro}$ from its primary amino acid sequence. The error and accuracy reports for generation of this structure are shown in Figure S7.

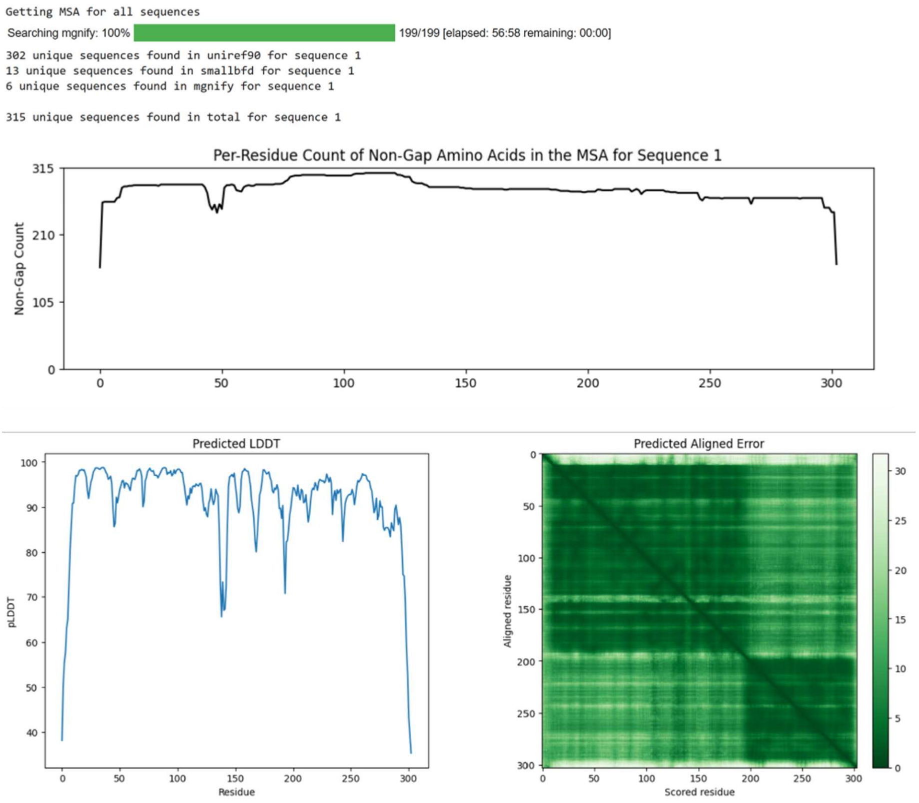


**Figure S7:** AlphaFold reports for the generation of a predicted crystal structure of HCoV-OC43 M$^{pro}$.

*S1.3.2 Generation of PL$^{pro}$ structures*

Predicted crystal structures for PL$^{pro}$ of OC43 and 229E were likewise generated using AlphaFold. Their error and accuracy reports are shown in Figures S8 and S9 respectively.

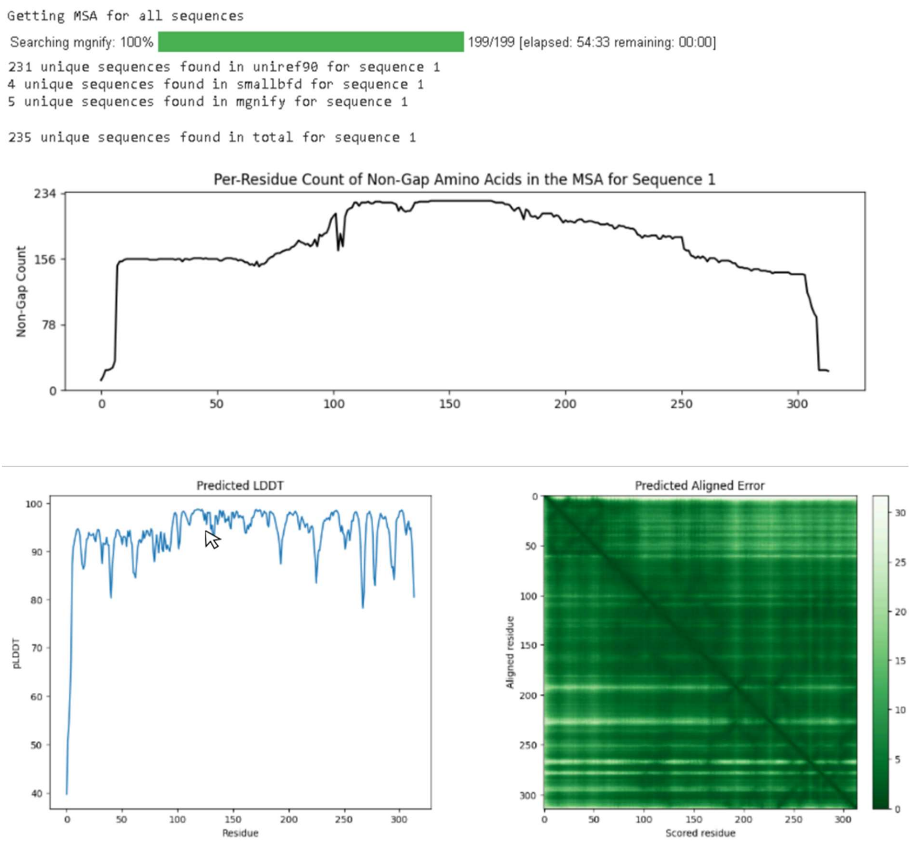


**Figure S8:** AlphaFold reports for the generation of a predicted crystal structure of HCoV-OC43 PL$^{pro}$.

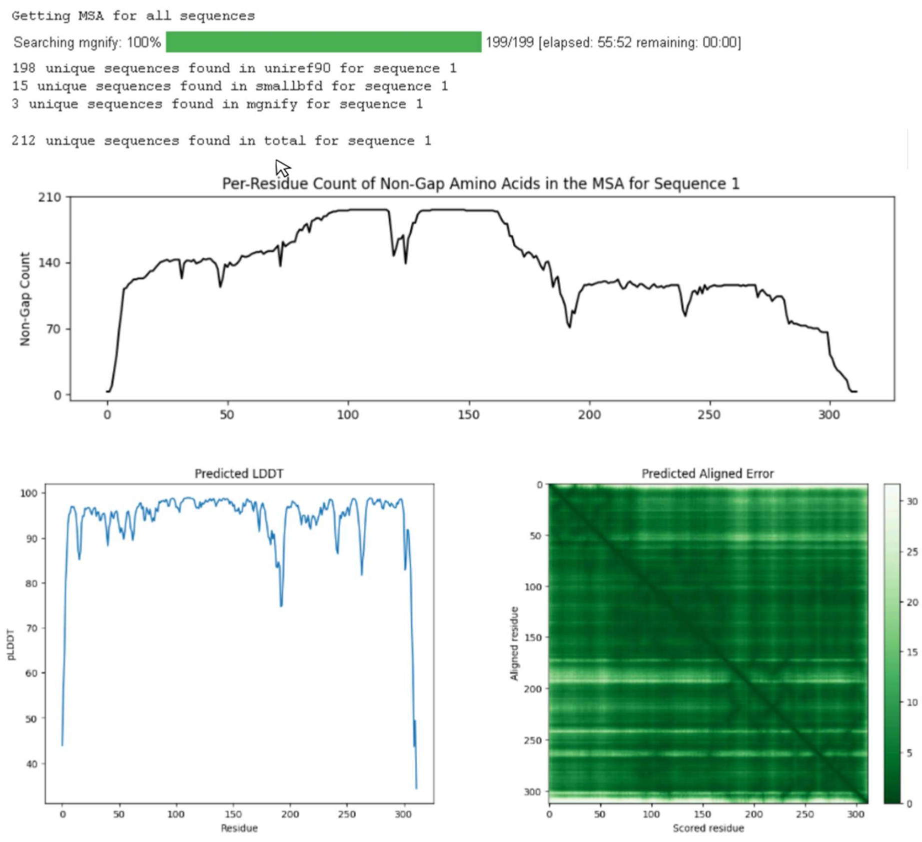


**Figure S9:** AlphaFold reports for the generation of a predicted crystal structure of HCoV-229E PL$^{pro}$.

### S2.0 EXPANDED RESULTS

#### S2.1 Enzyme Kinetics

Compounds **3**, **5**, **6**, and **7** showed *in vitro* inhibition of SARS-CoV-2 M$^{pro}$. Dose response curves were generated for all compounds that showed inhibition of enzyme protease

**Commented [CM1]:** not hyphenated…. in virology literature anyway.

activity (Figure S10). Only compound **5** showed *in-vitro* inhibition of SARS-CoV-2 PL$^{pro}$, and thus no other compounds' data is included here.

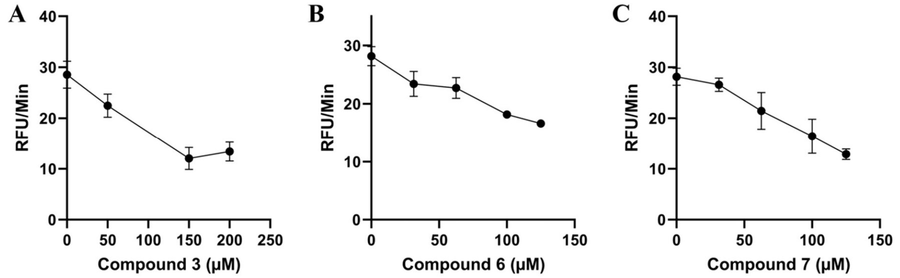


**Figure S10:** Dose response curves for less potent phenothiazine derivatives **3** (**A**), **6** (**B**), and **7** (**C**) assessed *in-vitro* with SARS-CoV-2 M$^{pro}$. Figures were generated using PRISM 9.4.0 (637) (https://www.graphpad.com).

#### S.2.2 *In-silico* Docking Studies

Compounds that showed *in-vitro* inhibition of protease activity were evaluated for their docking placement and interactions *in-silico*. Compounds **6** and **7** showed similar docking placements to one another (Figure S11 B, C). Their phenothiazine and adamantane moieties were in similar positions, and bonding interactions occurred with residue E166. Two π-hydrogen bonds formed between the NH of E166 and the center and lefthand phenothiazine rings of compound **6** (3.81 Å and 4.11 Å respectively), while a third formed between a CH of E166 and the center phenothiazine ring (4.02 Å). In the case of **7**, bonding occurred between the NH of E166 and the righthand phenothiazine ring (3.77 Å). Additional bonds occurred between the arene of H41 and a phenothiazine ring carbon of compound **7** (3.96 Å), a CH of M165 and the righthand ring (4.46 Å), and finally the carbonyl oxygen of Q189 and sulfur of **7** (4.06 Å). Compound **3** was observed in a docking configuration with its phenothiazine moiety pointed toward and bonding with N142 (Figure S11 A). Both the center and righthand rings of the phenothiazine formed π-hydrogen bonds with a CH of N142 (3.81 Å and 4.43 Å respectively).

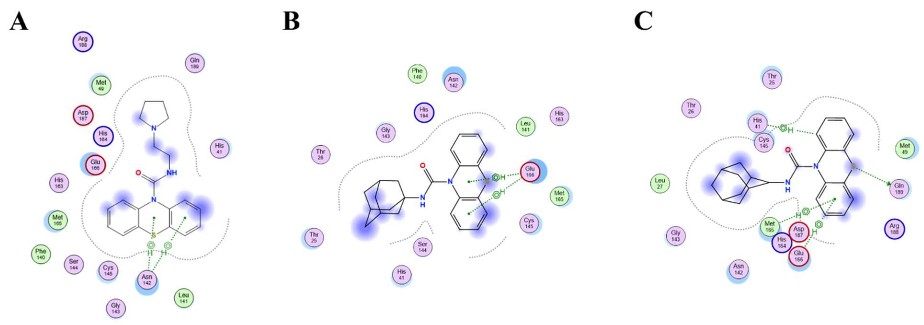


**Figure S11**: Ligand interaction maps for the binding of phenothiazine compounds **3** (**A**), **6** (**B**), and **7** (**C**) docked *in-silico* with SARS-CoV-2 M$^{pro}$ (PDB: 7L10). Docking and figures were generated with Molecular Operating Environment 2022.02 (https://chemcomp.com).

Top phenothiazine compound **5** was compared to the *in-silico* dock for nirmatrelvir with SARS-CoV-2 M$^{pro}$ for identification of potential similarities (Figure S12). Interactions between M$^{pro}$ and nirmatrelvir included two bonds between the CN warhead of nirmatrelvir and the SH and NH of C145 (2.17 Å and 2.56 Å respectively), as well as a third bond between the CN warhead and the NH of G143 (2.23 Å). Two other bonds were observed between a carbonyl oxygen of E166 and both an NH and carbonyl oxygen of the nirmatrelvir backbone (2.08 Å and 2.10 Å respectively). Finally, another bond between the lactone NH of nirmatrelvir and the terminal OH of E166 (1.91 Å) was observed. An additional intramolecular bond between a backbone NH and one of nirmatrelvir's tertiary methyl groups (2.32 Å) is also shown in Figure S12.

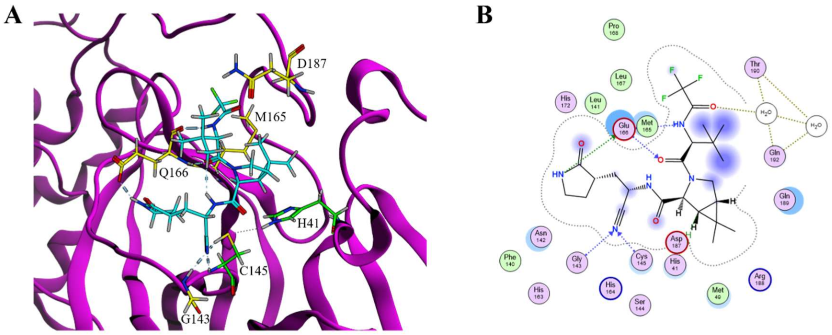


**Figure S12:** Top docking pose (**A**) and ligand interaction map (**B**) for nirmatrelvir with SARS-CoV-2 M$^{pro}$ (PDB: 7L10). Catalytic amino acids (H41 and C145) are shown with lime green carbons, other key amino acids are shown with yellow carbons. Figures were generated with Molecular Operating Environment 2022.02 (https://chemcomp.com).

### S2.3 Homology of M$^{pro}$ and PL$^{pro}$ Enzymes of SARS-CoV-2, HCoV-OC43, and HCoV-229E

#### *S2.3.1 Sequential Alignments*

Sequence alignments of M$^{pro}$ and PL$^{pro}$ enzymes of SARS-CoV-2, OC43, and 229E were done using Clustal Omega (https://www.ebi.ac.uk/Tools/msa/clustalo) and visualized with Jalview 2.11.2.7 (ttps://jalview.org) (Figure S13).

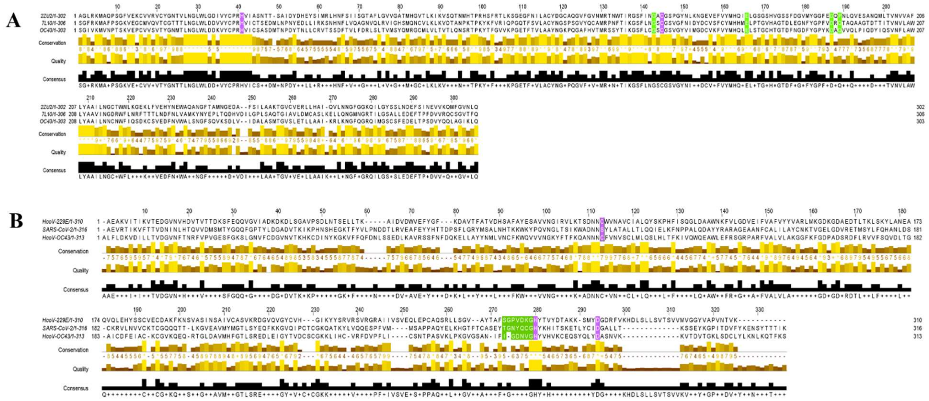

**Figure S13**: Primary sequence alignment for M$^{pro}$ (**A**) and PL$^{pro}$ (**B**) enzymes of SARS-CoV-2, HCoV-OC43, and HCoV-229E. Catalytic amino acids are highlighted in purple, other key amino acids are highlighted in green. Images were generated using Jalview 2.11.2.7 (https://jalview.org).

*S2.3.2 Structural Alignments*

Structural alignments of prepared enzyme PDB files for SARS-CoV-2, OC43, and 229E were done using MOE *Align/Superpose* feature. RMSD reports on the structural and sequential similarity among all three coronavirus enzymes are shown for M$^{pro}$ and PL$^{pro}$ in Figure S14.

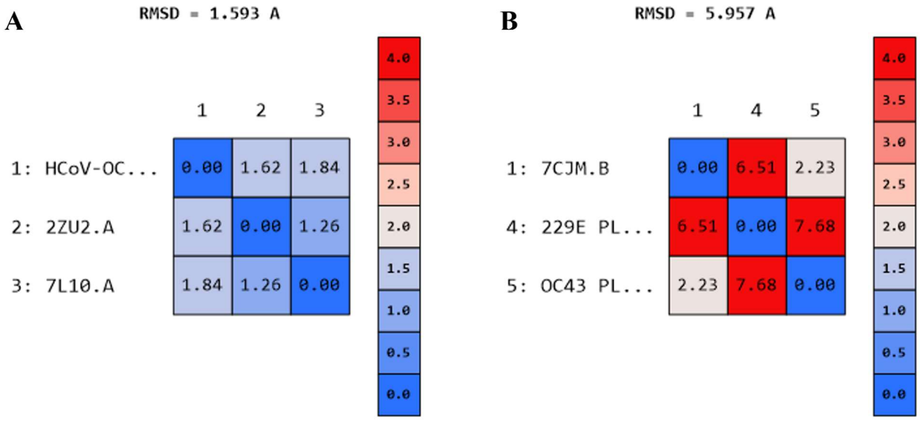


**Figure S14:** Root-mean square deviation report for alignment of M$^{pro}$ (**A**) and PL$^{pro}$ (**B**) enzymes of SARS-CoV-2 (M$^{pro}$: 7L10; PL$^{pro}$: 7CJM), HCoV-OC43 (AlphaFold PDBs), and HCoV-229E M$^{pro}$: 2ZU2; PL$^{pro}$: AlphaFold PDB). Alignments and figures were generated using Molecular Operating Environment 2022.02 (https://chemcomp.com).

#### S2.4 TCID$_{50}$ Assay

All phenothiazine compounds were evaluated in an HCoV-OC43 infection model to assess their potential to reduce viral titer. Assay results for compounds **1**-**4**, **6**, and **7** are shown in Figure S15.

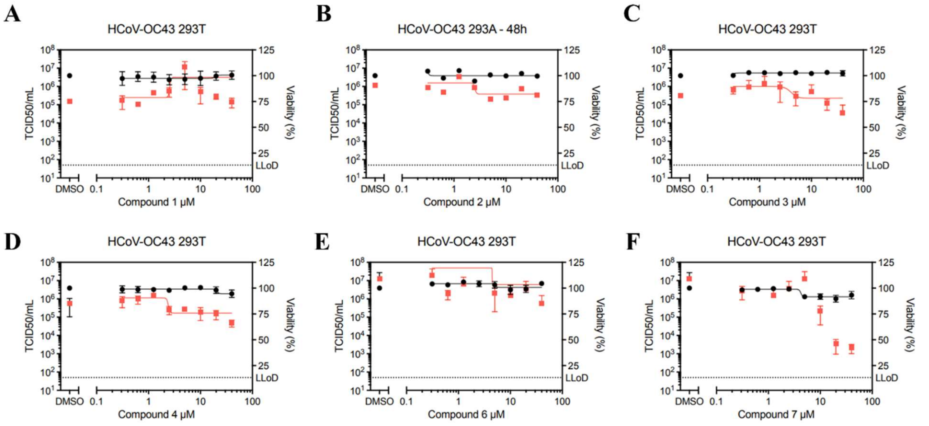


**Figure S15**: Reduction of viral titer by phenothiazine urea compounds **1** (**A**), **2** (**B**), **3** (**C**), **4** (**D**), **6** (**E**), and **7** (**F**)Viral titer was determined by 50% tissue culture infectious dose ($TCID_{50}$) assay and is shown as red squares. HEK293T cells were infected with HCoV-OC43 at an MOI of 0.1 then treated with 5 for 24 hrs before harvest and assayed with BHK cells. Percent cell viability of uninfected cells is shown as black circles and was measured in parallel. All experiments were completed in triplicate and their average values shown with standard error bars. Plots were generated using PRISM 9.4.0 (637) (https://graphpad.com).
